# Comprehensive and accelerated mapping of driver mutations through single-nucleotide random mutagenesis of target genes

**DOI:** 10.1101/2025.11.05.686667

**Authors:** Keisuke Yoshida, Elena Solovieva, Nami Miura, Shigeyuki Shichino, Michiaki Hamada, Masafumi Muratani, Takanori Amano, Kazufumi Honda

## Abstract

Identification of novel driver mutations is crucial for personalized medicine and drug discovery. However, genome-wide studies based solely on patient-derived data cannot confirm the oncogenic potential of detected mutations. Here, we present a genome editing-based method for comprehensive random mutagenesis of target genes. This approach generates cultured cells harboring random single-nucleotide variants (SNVs) within 10 days of DNA amplicon preparation. To validate its utility, random mutations were introduced into endogenous exons of lung cancer-related oncogenes in HEK293 cells. Spheroid formation assays and xenograft models demonstrated enhanced tumorigenicity, indicating the acquisition of oncogenic potential. Amplicon-sequencing of spheroid transcripts revealed significant enrichment of variants annotated as pathogenic in clinical databases and computational predictions. Furthermore, osimertinib treatment of lung cancer cell lines with random SNVs on EGFR identified 58 novel resistance mutations. These results highlight the effectiveness of our mutagenesis platform, offering a powerful tool to advance cancer research and precision medicine.

## Introduction

The identification of cancer-associated mutations through genome-wide studies of clinical samples has greatly contributed to the development of optimized therapeutic strategies, improving our understanding of the mechanism of resistance against conventional molecular-targeted therapies, and facilitating the development of novel molecular-targeted drugs^1,2^. However, genomic studies of patient-derived samples require substantial time, financial resources, and labor. Obtaining sufficient specimens for a reliable analysis is challenging, especially for diseases with a limited number of cases such as rare cancers^3,4^. In addition, although a typical tumor genome harbors thousands of somatic mutations, a small subset are driver mutations, and the vast majority are biologically neutral passenger mutations^5,6^. Distinguishing true drivers from passengers in such mutation-heavy backgrounds remains a major challenge in cancer genomics^7,8^. Therefore, in addition to “top-down” cancer genome analyses using clinical specimens, experimental methods evaluating the oncogenic potential of mutations, representing a “bottom-up” approach by cell-based assays, need to be designed to identify and comprehensively understand driver mutations.

High-throughput CRISPR/Cas9 saturation mutagenesis screening enables large-scale genetic perturbations at a genome-wide scale^9,10,11,12^. Genome-wide single guide RNA (sgRNA) libraries can be deployed to disrupt or modify target genes, which facilitates the systematic identification of essential genes and functional variants in their native contexts^13,14,15^. These approaches are used for screening novel functions of genes and variants including drug resistance and for predicting potential underlying mechanisms. For example, saturation editing^9^ uses overlapping sRNAs to induce diverse mutations in a target gene, thereby identifying critical protein domains and specific mutations that drive therapeutic resistance. Despite the potential of these high-throughput genome editing screening system for improving our understanding of variant function in cancer genome^16^, the preparation of gRNA library targeting specific genome region requires extensive time and labor, which limits the execution of this type of experiment. In this study, we propose an experimental system for the comprehensive identification of functional variants on target genes via the rapid and simple construction of a gRNA library.

## Results

### Genomic region-specific mutagenesis

The aim of this study was to establish cellular phenotype-based rapid screening of driver mutations on target genes as bottom-up cancer genome analysis. To achieve this goal, random single-nucleotide mutations were introduced into target genomic regions in cultured cells by combining base editor (BE) enzyme technology with a gRNA expression vector library (**Fig. 1a**). To prepare a gRNA library from a DNA fragment of the target genomic region, PCR amplicon was first randomly fragmented (Extended Data Fig. 1). A stem-loop adaptor containing the recognition site for a TYPE-III restriction enzyme was introduced into the terminals of the fragmented target DNA, followed by treatment with restriction enzymes, which theoretically should lead to the production of a 20-bp DNA fragment as the spacer sequence of gRNA. The 20-bp spacer with transactivating CRISPR RNA (tracrRNA) sequences was inserted into a U6 promoter-driven sgRNA expression *E.coli* vector, which allows the preparation of a gRNA library with a spacer sequence containing 20 bp of the target sequence within 3 days (Extended Data Fig. 1). A near protospacer adjacent motif (PAM)-less BE^17^, in which a highly efficient PAM is NRN, can induce point mutation at 5–7 nt from the 5’ terminal of the spacer sequence. Co-transfection of near PAM-less BE and gRNA library expression vectors into cultured cells was expected to randomly introduce point mutations into endogenous target genomic regions because different gRNA vectors targeting different sites were incorporated into each single cell. In the present experimental scheme, if a target DNA amplicon is already prepared, a cellular population harboring random mutations in target genomic regions can be obtained within 10 days (**Fig. 1a**).

**Figure 1.**
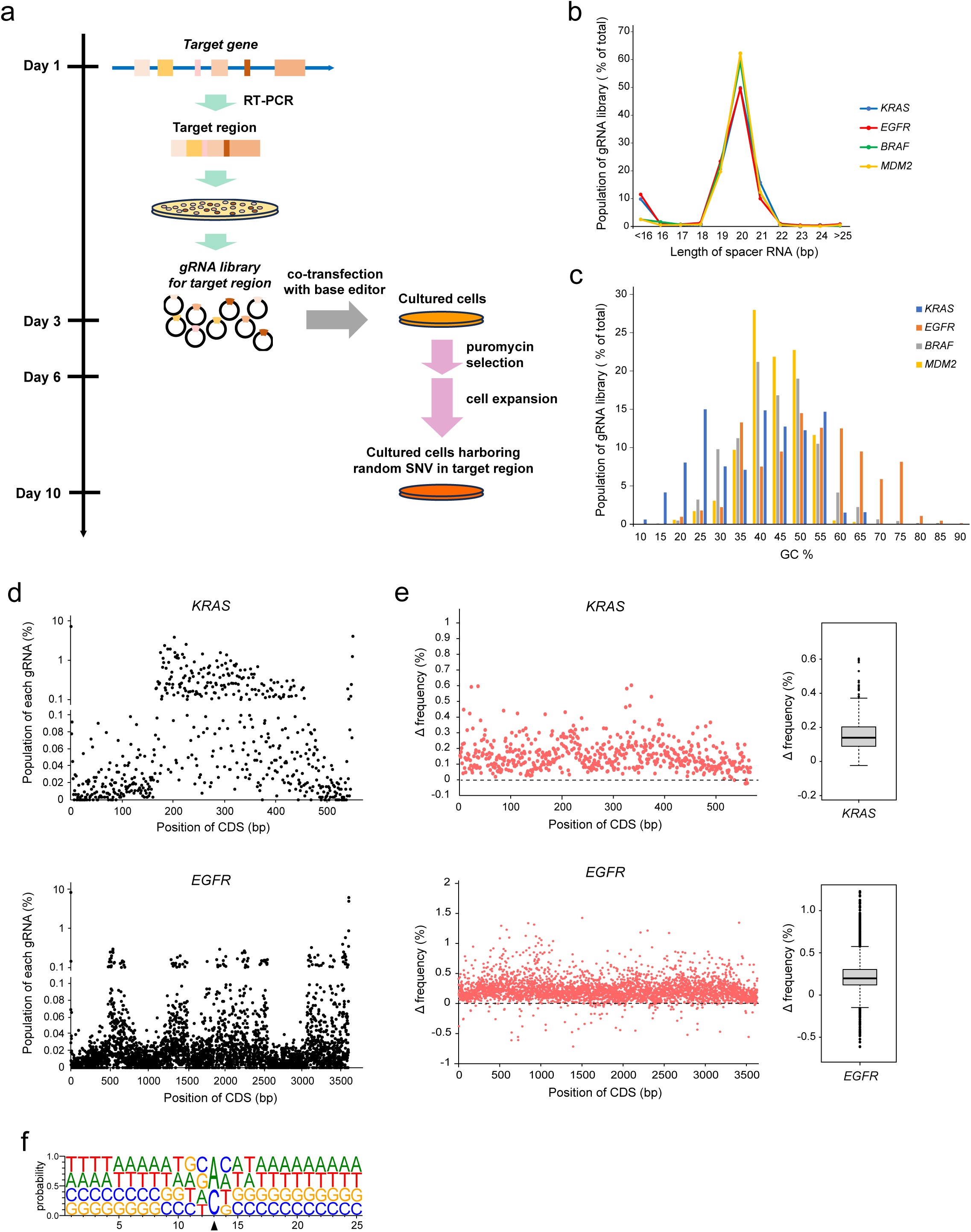
Random mutagenesis of target genomic regions achieved through rapid construction of a gRNA library. **a**, Experimental scheme for the introduction of random single-nucleotide variants (SNVs) into the target genomic region. **b**,**c**, Length (b) and GC content (c) of the spacer sequence in gRNA libraries. A DNA fragment of the exon sequence from four genes was converted into a gRNA library. **d**, Proportion of gRNA in which the spacer sequence corresponds to 20 bp along the exon sequence of KRAS (top) or EGFR (bottom). **e**, Frequency of SNVs on all bases of the KRAS (top) or EGFR (bottom) gene. The frequency distribution is shown on the right (n = 552 for KRAS, n = 3,615 for EGFR) by box plot. HEK293 cells were treated with BEs (control) or both BEs and gRNA library (treatment), and mutation frequency was assessed in samples collected after puromycin selection with timing corresponding to day 6 in a. Change in frequency was determined by subtracting control frequency from treatment frequency. f, Probability of nucleotide appearance at 25 bp around the induction site. The site of observed SNVs with >0.05% Δfrequency (n = 21,988) is adjusted according to the center position and indicated by a black triangle. For box plots in e, elements indicate the following values: center line, median; box limits, upper and lower quartiles 1.5× interquartile range; and points, outliers.

To verify the proposed experimental strategy, we first prepared a gRNA library for the exon regions of six oncogenes for which driver mutations are abundantly detected in lung cancer patients^18^, namely *KRAS*, *BRAF*, *EGFR*, *CDK4*, *MDM2,* and *MET*. PCR amplicons of protein coding sequences (CDS) of these genes were prepared by RT-PCR using cDNA collected from HEK293 cells and used as starting material for the assay. In the gRNA library prepared using this method, 82.9–94.3 % of the gRNA vectors had a spacer sequence measuring 19–21bp in length, which is compatible with spCas9^19^ (**Fig. 1b**). The GC content of the spacer sequence in the majority of vectors was 40–55%, which is within the desirable range for functional gRNA^20^ (**Fig. 1c**). We mapped the spacer sequences in the target genomic sequence and found that they were mostly distributed evenly along the target DNA sequence and most of them were aligned without a gap (**Fig. 1d**; Extended Data Fig. 2a,b). To estimate the off-target effect of the libraries, 20-bp window-size sequences of target regions were mapped into the whole genome sequence. Off-target regions were identified in fraction of gRNAs (Extended Data Fig. 2c). Classification of the off-target regions showed that >97.9% of the gRNAs in *KRAS*, *BRAF*, and *MDM2* libraries corresponded to non-coding regions (Extended Data Fig. 2d). For *EGFR*, 11 spacer sequences had off-target coding regions of *EGFR* homologous genes such as *ERBB2/3*, suggesting that induction efficiency in some on-target sites could be inhibited by the dispersed recruitment of the BE to off-target sites. These results indicated that the gRNAs included in these vector libraries predominantly targeted the intended gene loci, especially within coding regions.

Next, to investigate the mutation frequency profile induced by the experimental system on target genomic regions, the gRNA libraries were co-transfected with the expression vectors of cytidine base editor (CBE), CBEmax-SpRY-Puro and adenine base editor (ABE), ABEmax-SpRY-Puro, into HEK293 cells; these cells were derived from normal healthy human embryonic kidney and had no driver mutations in the six genes at least^21^. The single-nucleotide mutations were introduced in a dense pattern along target exon regions (**Fig. 1e,f**, Extended Data Fig. 3a). Unexpectedly, not only transition mutations, but also transversion mutations were induced, and certain types of transversion mutations were dominantly detected in variants at lower induction levels (Extended Data Fig. 3b,c). The BE enzymes were reported to induce transversion mutations with lower efficiency^22^, and enzymatic interaction between ABE and CBE might potentiate this side-effect. Besides the single-nucleotide replacements, insertions and deletions which the BE does not induce, were not detected (Extended Data Fig. 3d). The results suggest that the experimental system could induce random mutations into targeting-endogenous genomic regions at least for coding regions. The protocol was thus named BELT (<u>B</u>ase editor and <u>E</u>nzymatically prepared gRNA <u>L</u>ibrary-based gene locus-driven <u>T</u>argeting) mutagenesis.

### BELT mutagenesis of oncogenes

To assess the effect of BELT mutagenesis on oncogenicity, HEK293 cells harboring single-nucleotide random mutations on six oncogenes were transplanted into immunodeficient mice. Tumor formation was higher in this cell fraction than in control HEK293 cells (**Fig. 2a-c**: Extended Data Fig. 4a). Anchorage-independent growth is a cancer-specific phenotype, and assessment of this cellular phenotype is traditionally used to evaluate oncogenicity^23,24^. BELT-treated cells cultured on an ultra-low attachment dish showed strong ability of spheroid formation (Extended Data Fig. 4b), indicating that BELT mutagenesis on oncogenes resulted in the acquisition of oncogenic potential.

**Figure 2.**
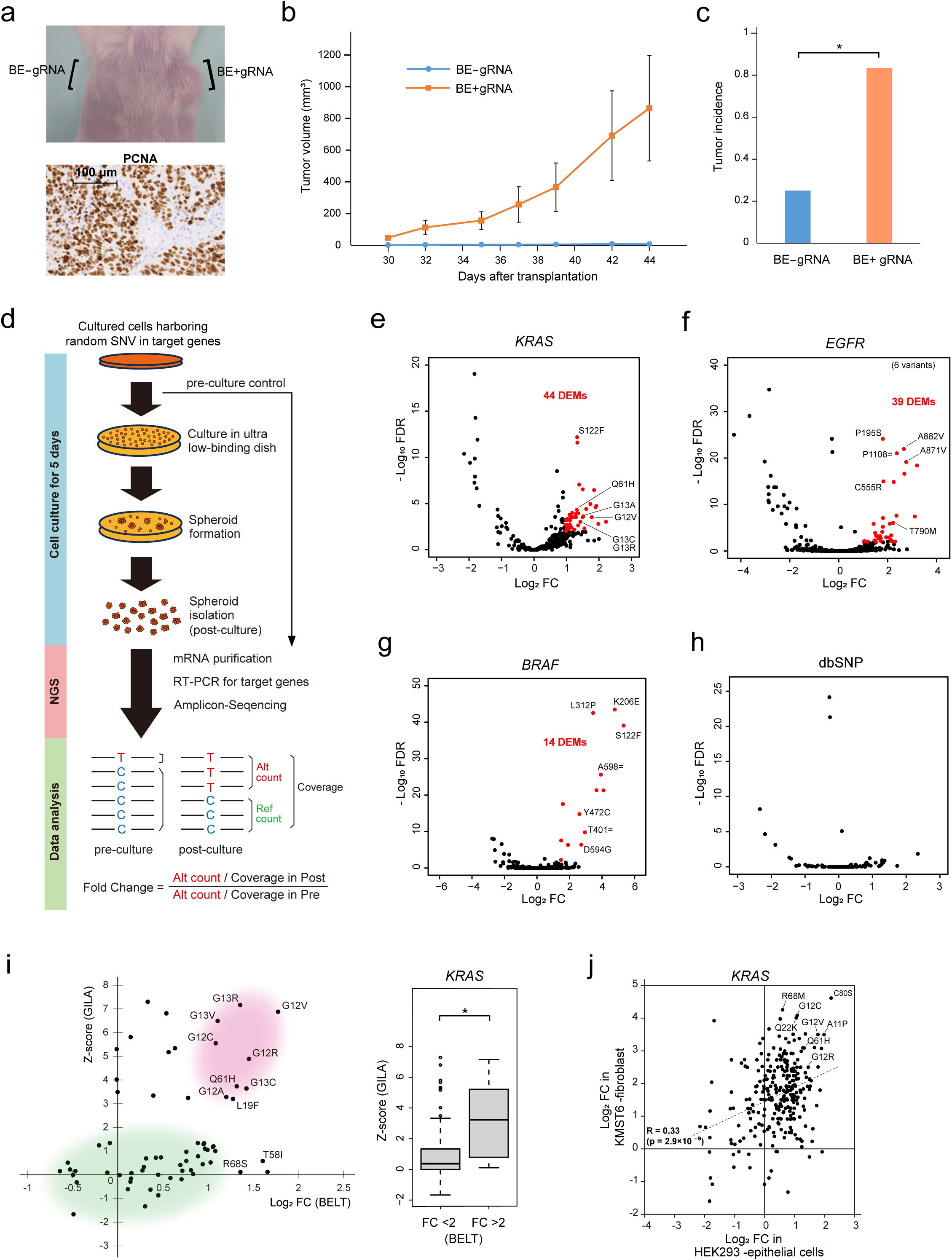
Specific detection of known driver mutations by BELT screening. **a**, Image of the shoulder of an SCID mouse inoculated with HEK293 suspension (top). HEK293 cells treated with only BEs or both BEs and gRNA library for the coding region of KRAS, BRAF, EGFR, CDK4, MDM2, and MET were transplanted into the left or right shoulder, respectively. Representative image of PCNA staining of a cell mass formed in vivo (bottom). **b**, Change of tumor volume 30–44 days after transplantation (n = 12). Values represent the average ± SE. **c**, Tumor incidence from cells with or without induction of random SNVs in six oncogenes (n = 12). P value was calculated by Fisher’s exact test. **d**. Experimental scheme describing the screening for driver mutations by BELT mutagenesis. **e-h**, Volcano plot of the profiles of COSMIC variants on KRAS (e), EGFR (f), and BRAF (g), and of dbSNP variants on six oncogenes (h) in HEK293 cells. Each dot indicates a variant. The horizonal axis indicates the FC of mutation frequency between pre- and post-culture in a low binding dish, and the vertical axis indicates FDR reflecting the statistical significance of the difference (Fisher’s exact test). DEMs (FC >2, FDR <0.01) are colored in red, and representative mutations are shown with amino acid alterations. The number of data points falling outside the plotting range is shown in the plot margins. **i**, Scatter plot for KRAS mutations with FC between the current results and previously reported data (left). Areas of densely populated variants with higher or lower FC in both datasets are indicated in red or green, respectively. The box plot indicates the distribution of z-scores from previous data divided into FC >2 (n = 16) and FC <2 (n = 54) compared with the present results (right). P value was calculated by Wilcoxon rank-sum test. j, Scatter plot of mutations with FC in the KRAS gene between epithelial-like and fibroblast-like cell lines in the present study. *p < 0.05.

We expected that cancer-supporting driver mutations were enriched in spheroids from BELT-treated HEK293 cells cultured on ultra-low attachment dish. To verify this possibility, mutation frequency on target genes was analyzed using NGS by comparing cell fractions from pre- and post-culture on ultra-low attachment dish **(Fig. 2d)**. Following to this analysis for these two samples, fold-change (FC) and p value in cancer-associated SNVs registered in the COSMIC cancer mutation database were calculated for the CDSs of six target genes. To confirm the statistical significance of this calculation, the probability of mutation enrichment was estimated by Bayesian inference for the read count per each coverage with the prior weight of the quality score. The results showed that all variants filtered using the threshold as FC> 2.0 and, FDR< 0.01 (Fisher’s exact test) had >0.99 probability, supporting the statistical reliability of the candidate driver mutation selection (Extended Data Fig. 4c). Differentially enriched mutations (DEMs) by anchorage-independent culture were identified using these criteria (**Fig. 2e-g**, Extended Data Fig. 4d-f and Supplementary Table S1). In *KRAS*, the top three hot spots, G12X, G13X and Q61X in *KRAS*-positive cancer patients^25^, were included among DEMs (**Fig. 2e**). COSMIC mutations experimentally certified as cancer drivers such as EGFR T790M^26^ and BRAF D594G^27^ were also found in DEMs (**Fig. 2f,g**). On the other hand, DEMs were not detected in variants registered in dbSNP, most of which are recognized as non-pathogenic (**Fig. 2h**).

To verify the reliability of the BELT screening results for driver mutations, each representative DEM that was not registered as a pathogenic mutation in OncoKB and ClinVar databases was introduced into HEK293 cells. Cell lines harboring these SNVs showed increasing of anchorage-independent growth (Extended Data Fig. 4g and Extended Data Table 1). When our dataset was compared with previous result^12^ in COSMIC mutations on *KRAS*, variants in hot spots with higher scores were concordantly observed in both datasets (**Fig. 2i**). We also performed BELT screening in the human fibroblast cell line KMST6, and hotspot mutations such as KRAS G12X and BRAF around codon 600 showed a higher FC in both cell-lines (**Fig. 2j**, Extended Data Fig. 4h and Supplementary Table S1). These results indicate that BELT screening for driver mutation in target genes has high utility and sufficient reliability.

### Functional feature in DEMs

Next, we compared our dataset with public tumor databases such as OncoKB and ClinVar. In OncoKB, DEMs detected in *KRAS* variants from the COSMIC database only include pathogenic and likely pathogenic variants (**Figure 3A**). ClinVar variants include DEMs in six genes, and their annotations were analyzed. Pathogenic and likely pathogenic variants were significantly enriched in DEMs, whereas benign and likely benign variants were not enriched (**Figure 3B** and Extended Data Fig. 5a). The annotation according to disease showed that variants associated with lung cancer were highly enriched in DEM (Extended Data Fig. 5b).

**Figure 3.**
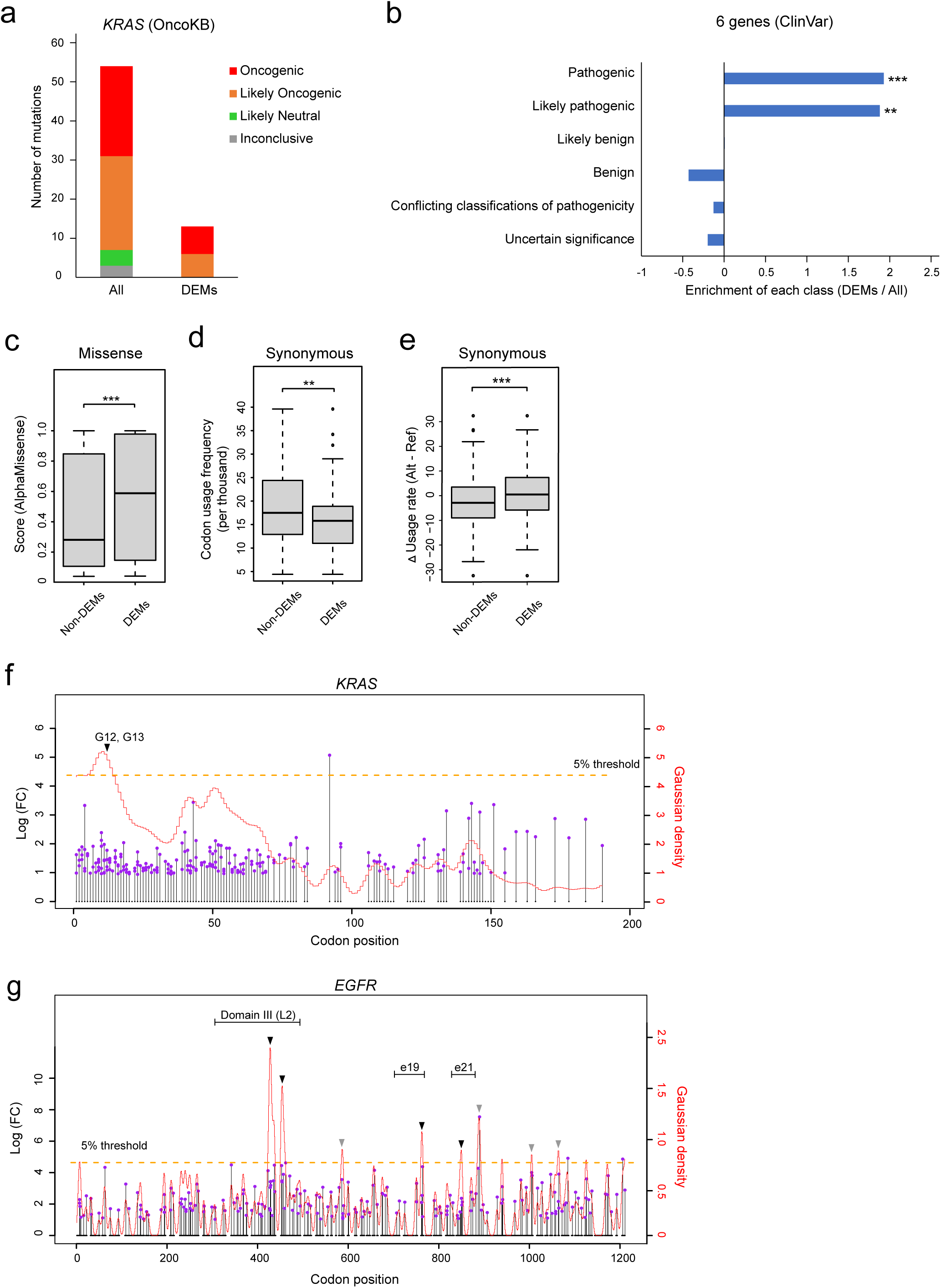
Clinical characteristics of the mutation cohort identified through experimental methods. **a**, Stacked bar plot of the number of mutations classified according to pathogenic annotations in OncoKB. DEMs were separated from the list of COSMIC mutations on KRAS and analyzed. **b**, Enrichment of the variant population with annotations of clinical significance in ClinVar between DEMs (FC >2, FDR <0.01) and all mutations. P value was calculated by Fisher’s exact test. **c**, Distribution of AlphaMissense scores of missense variants between non-DEMs (n = 6,810) and DEMs (n = 166). P value was calculated by Wilcoxon rank-sum test. **d,e**, Distribution of usage frequency in the reference codon (d) and subtraction of the reference from the alteration frequency with SNVs (e) between non-DEMs (n = 2,865) and DEMs (n = 69). P values were calculated by Wilcoxon rank-sum test. **f,g**, Lollipop plot of DEMs (FC >2, FDR <0.05) with FC at 1 bp resolution along KRAS (f) and EGFR (g) in HEK293 cells. The Gaussian density curve of FC is shown as a red line, and the top 5% density region is shown as an orange dot line. Hot spots of the density of DEMs above the 5% threshold are indicated by gray triangles, and those overlapping with clinical hotspots and the Receptor L domain of EGFR are shown in black triangles. For box plots in c-e, elements indicate the following values: center line, median; box limits, upper and lower quartiles 1.5× interquartile range; and points, outliers. *p < 0.05, **p < 0.01, ***p < 0.001.

The functional features of mutations were then analyzed by combining DEMs detected in the COSMIC and ClinVar databases. Among the DEMs in these variants, 69.0% were missense and 28.2% were synonymous mutations (Extended Data Fig. 5c). The transition mutations showed higher enrichment in DEMs than in non-DEMs, indicating the pathogenic tendency of transition mutations (Extended Data Fig. 5d). For missense mutations, the effect of a single amino-acid substitution was assessed using scores obtained from computational prediction databases that evaluate the impact of genetic variants on protein function. In PolyPhen-2^29^, which predicts the effect of amino acid changes on protein function rather than on disease, functional scores were similar between DEMs and non-DEMs (Extended Data Fig. 5e). On the other hand, CADD^30^, ClinPred^31^, and especially AlphaMissense^32^ databases showed enrichment of higher score in DEMs than in non-DEMs (**Fig. 3c** and Extended Data Fig. 5e). BLOSUM62 matrix^33^ which predicts the impact of amino acid substitutions, also showed a lower score in DEMs, likely indicating the effect of non-conservative amino acid change on protein structure or function (Extended Data Fig. 5f).

We analyzed the characteristics of synonymous mutations in DEMs to understand the molecular mechanisms by which synonymous mutations contribute to oncogenic potential. Three known molecular mechanisms underlie the effect of synonymous mutations on protein function^34^: alterations in splicing regulation, changes in mRNA secondary structure, and codon usage bias. The effect on splicing regulation was investigated by calculating SpliceAI^35^ scores for the identified synonymous DEMs, which showed that scores were close to zero, indicating minimal predicted impact on splicing (Extended Data Fig. 6a). The effect of synonymous DEMs on minimum free energy and base paring probability of the predicted mRNA secondary structure was investigated using the ViennaRNA-based tool^36^, which showed similar scores between DEMs and non-DEMs (Extended Data Fig. 6b,c).

Analysis of usage frequency in reference codons with synonymous DEMs showed that it was significantly lower in DEMs than non-DEMs (**Fig. 3d**). When the reference codon was substituted by an altered codon, the codon usage frequency of the altered codon was observed to be higher in DEMs than in non-DEMs (**Fig. 3d** and Extended Data Fig. 6d). Furthermore, uracil was enriched in the third position of the reference codon in synonymous DEMs, and the codon for cysteine, which forms disulfide bonds that are important for proper protein folding^37^, was enriched as well (Extended Data Fig. 6e,f). These observations support the presence of codon bias in a portion of synonymous DEMs, which might have functional relevance regarding the efficiency of protein translation.

To investigate the distribution of driver mutations along the coding sequence of oncogenes, DEMs were identified in the target gene regions at single-nucleotide resolution, including known mutations and those not yet reported in human clinical samples. The lollipop plot indicates that DEMs were enriched in the N-terminal of the *KRAS* coding region; the region with the highest mutation density corresponded to the G12/13 codon (**Fig. 3f**; Supplementary Table S2). Eight DEMs hotspots were identified in the *EGFR* gene; the genomic domain of exons 19 and 21, which contained the hotspots of driver mutations such as exon 19 deletion and L858R, overlapped with the corresponding experimentally identified hotspot (**Fig. 3g**; Supplementary Table S2). These results support the reliability of driver mutations identified by BELT screening and provide novel insight into driver mutations that can enhance our understanding of the molecular mechanisms underlying oncogenesis.

### Rapid detection of drug-resistant SNVs

Next, we used BELT screening to identify mutations associated with resistance to molecular-targeted drug. Osimertinib is a third-generation EGFR tyrosine kinase inhibitor (TKI), that is currently the first-line treatment of *EGFR*-positive lung cancer^37^. Random mutations were introduced into the exon regions of the *EGFR* gene in a cell line carrying a hotspot of driver mutation, L858R and exon19 deletion. Mutations associated with resistance to osimertinib were enriched through continuous culture in a medium containing the drug (**Fig. 4a**). In the human lung adenocarcinoma cell line II-18 or PC-9 harboring a heterozygous L858R or homozygous exon19 deletion (E746–A750del) respectively, comparison between osimertinib- and DMSO-treated samples identified 54 DEMs and 8 DEMs as osimertinib-resistant mutations (**Fig. 4b,c** and Supplementary Table S3). Regarding resistant mutations in the two cell lines, half of the DEMs in PC-9 overlapped with those in II-18 (**Fig. 4d**). To verify the functional effect of mutations identified by the drug resistance screening, a representative DEM for each gene was introduced into II-18 cells. All nine mutations increased resistance against osimertinib (**Fig. 4e**). Assessment of the dynamics of EGFR activity during osimertinib treatment showed that II-18 cells carrying the M600V, V592=, or I744V mutation on *EGFR* exhibited persistence of EGFR phosphorylation in response to the drug, indicating that the inhibition caused by these mutations was impaired (**Fig. 4f**). To investigate the functional interaction of mutations on the *EGFR* allele, the pairing patterns were analyzed. The results showed that 46 mutation pairs were significantly enriched in osimertinib-treated sample relative to the DMSO control, and L858R acted as a central hub (**Fig. 4g**). In certain mutations, drug resistance is associated with factors cis- or trans-interacting with driver mutation^38^; we thus investigated variants enriched after the separation of reads with or without the L858R mutation (Extended Data Fig. 7a,b). The results indicate that certain mutations were specifically detected in the L858R allele, supporting the cis-effect of resistance on osimertinib (Extended Data Fig. 7c). Mapping of resistant mutations at 1-bp resolution identified eight mutation hotspots, which partly overlapped with driver mutation hotspots (Extended Data Fig. 7d). This suggests that a portion of resistant mutation also had driver activity, such as T790M^26^.

**Figure 4.**
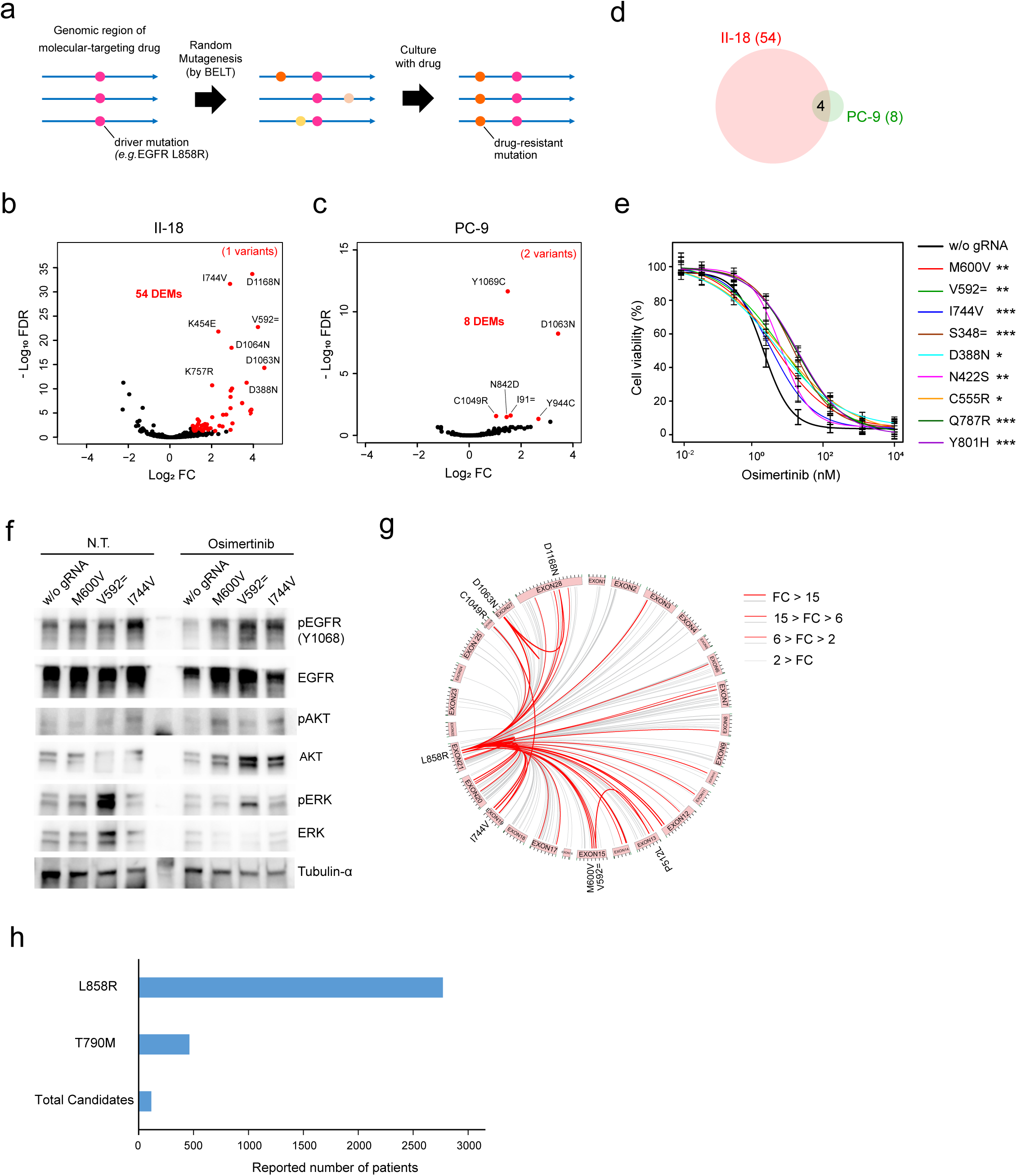
Identification of osimertinib-resistant SNVs by BELT screening. **a**, Model of drug-resistant mutation enrichment. **b,c**, Volcano plot for COSMIC variants on EGFR in II-18 (b) and PC-9 (c) cells. Each dot indicates a variant. The horizonal axis indicates the FC of mutation frequency between DMSO and osimertinib treatment in a low binding dish, and the vertical axis shows FDR reflecting the statistical significance of the difference (Fisher’s exact test). DEMs (FC >2, FDR <0.05) are colored in red, and representative mutations are shown with amino acid alterations. P values were calculated by Fisher’s exact test. The number of data points falling outside the plot range is annotated in the plot margins. **d**, Venn diagram of DEMs between II-18 and PC-9 cells. **e**, Response to different doses of osimertinib in different II-18 cells. To estimate the difference of IC50 values between wild-type and mutant lines, P values were calculated by the Wald test by comparing the estimated ED values, based on the standard error of the difference and the standard normal distribution. **f**, The total and phosphorylated levels of EGFR, AKT, and ERK proteins in II-18 cells treated with or without osimertinib for 2 h. **g**, The circos plot visualizing SNV pairs in the EGFR gene. Pink boxes denote CDS regions with chromosomal positions. Links connect pairs of two SNVs located in the same read and position. Mutation pairs with counts >10 in osimertinib samples are shown, and the associations of significantly enriched pairs in osimertinib relative to DMSO samples (FC >2, FDR <0.05) are indicated in red. Line thickness corresponds to FC as indicated in the annotation on the right side of the figure. The heatmap represents the number of SNV pairs containing the SNV at a given position. P values were calculated by Fisher’s exact test. **h**, Reported number of patients. The numbers of cases were obtained from the COSMIC database, and the total number of patients was calculated as the sum of cases with osimertinib-resistant SNVs identified in this study. For dose responsive curve in e, it was prepared by fitting the data points to a four-parameter logistic model and the error bars represent the confidence intervals of the predicted response values at each dose level based on the fitted model.

Here, we identified 58 *EGFR* mutations potentially associated with osimertinib resistance. Although each mutation has been reported only a few times individually (Extended Data Fig. 7e), the cumulative number of reports was approximately 25% of those for the T790M mutation, which is the most frequently observed resistance mutation in cases resistant to first- and second-generation EGFR-TKIs^39^ (**Fig. 4h**). These results indicate the utility of BELT screening for variants related to therapeutic effect of a drug.

## Discussion

In this study, we established BELT screening, a novel genome editing-based method for comprehensive random mutagenesis of target genes. The driver mutations detected by this method through unbiased selection of mutation sites were enriched in pathogenic variants certified epidemiologically and experimentally. Compared with previous studies describing CRISPR-based screening of functional variants^9,10,11,12,13^, the method developed here markedly decreases the time and cost needed for the procedure while maintaining single-nucleotide resolution because the gRNA library derived from *E.coli* can be readily constructed using standard cloning techniques (Extended Data Table 2). Experimental identification of pathogenic variants as bottom-up cancer genomics could be of value to analyze orphan diseases including cancers with a small number of cases. Excluding missense mutations, computational prediction of driver mutations in non-coding regions such as introns, untranslated mRNA, and gene promoters is difficult because these mutations do not change the amino acid sequence and do not directly alter protein conformation. In BELT screening, the target genomic region can be freely selected using a PCR amplicon as starting material DNA for gRNA library preparation (Extended Data Fig. 1), thereby allowing functional evaluation of variants of uncertain significance in exon regions as well as in non-coding regions. Mutations related to spheroid formation or drug resistance were identified in this study using different cell lines, suggesting the utility of the method for exploring cell type-specific driver mutations and functional interactions of pathogenic mutations. The driver mutations identified in the study correlated with pathogenic scores in computational prediction tool such as AlphaMissense^31^. This dataset could also serve as training data for artificial intelligence models predicting protein function.

In this study, we used the method developed to systematically evaluate the driver potential of single-nucleotide substitutions across whole exons of *KRAS* and *EGFR* oncogenes at 1-bp resolution. Importantly, we found that regions exhibiting peaks in functional driver activity overlapped precisely with mutational hotspots observed in clinical tumor samples. This concordance strongly supports the notion that these hotspot mutations are not merely frequent by chance, but are under positive selection due to their functional importance in oncogenesis. Furthermore, the present findings validate the underlying assumption of our approach, namely, that mutations promoting anchorage-independent growth *in vitro* are indicative of functional contributions to tumorigenesis *in vivo*. The ability of this system to recapitulate clinically relevant mutation patterns underscores its utility for the functional annotation of cancer variants and highlights its potential for identifying both established and previously unrecognized driver mutations with oncogenic potential.

The annotation results from cancer genome databases support the reliability of the driver mutations identified in our analysis. However, in the gRNA library generated using this method, a minor population of gRNAs may bind to unintended sites in addition to target genes, potentially resulting in off-target effects. Although the likelihood is considered low, if mutations introduced by off-target activity possess driver-like function, there is a risk that a passenger mutation in cells harboring that mutation might be erroneously enriched and misclassified as a driver mutation. To avoid such effects, it is essential to design highly-specific gRNAs and to synthesize each gRNA individually, as described previously^11,12,13^. Because the BELT method allows for the introduction of only single-nucleotide substitutions, comprehensive incorporation of other types of mutations, such as short InDels and multi-nucleotide variants, would require alternative approaches. One such strategy is the use of donor oligonucleotides combined with homologous recombination, as exemplified by saturation genome editing^9,13^.

Among the driver mutations identified in six oncogenes, we observed a notable enrichment of transition mutations. This finding is consistent with previous reports and suggests a potential bias in the mutational processes underlying oncogenesis. Compared to transversions, missense mutations arising from transitions tend to substitute amino acids with similar physicochemical properties^40,41^. Such conservative changes may allow the mutant proteins to retain their structural integrity while acquiring gain-of-function characteristics. This notion is further supported by the observation that missense driver mutations often exhibit lower BLOSUM62 scores^32^, which is indicative of non-conservative substitutions. Collectively, these results suggest that transition mutations could provide an evolutionary advantageous route to tumorigenesis by enabling functionally beneficial changes while minimizing detrimental effects on protein stability.

We found that synonymous DEMs were likely to be detected in codons with relatively low usage frequency, and notably, the mutations often resulted in a shift toward codons with higher usage. Synonymous mutations that increase or decrease codon optimality can enhance or suppress translational efficiency, respectively^42^. Moreover, local changes in translation kinetics can influence co-translational protein folding, thereby affecting the final protein structure^43,44^. Consistent with this, we observed an enrichment of codons encoding cysteine residues that participate in disulfide bond formation, a structural feature critical for tertiary protein folding^36^. Additionally, synonymous driver mutations showed a preference for codons ending with uracil (U) at the third nucleotide position. This raises the possibility that these mutations interact functionally with wobble-base pairing mechanisms^45^, particularly those involving cancer cell–specific modified tRNAs such as queuosine^46^, thereby modulating translation efficiency.

Although our findings suggest that these synonymous driver mutations may contribute to increased protein expression or altered protein structure in oncogenes, we cannot exclude possibility that a portion of synonymous DEMs contributes to mRNA secondary structure changes. For example, one of the synonymous mutations in our dataset, KRAS G10=, increases protein expression by altering mRNA secondary structure^47^. However, these observations are based on the analysis of only six oncogenes and may reflect gene-specific trends. The generalizability of these findings to other oncogenes or tumor suppressor genes remains to be determined. Indeed, *KRAS* contains a high proportion of rare codons^48^, which may make it particularly susceptible to changes in translation efficiency caused by synonymous mutations.

In our analysis of resistance mutations to EGFR-targeting drug, the number of mutations detected in the II-18 cell line was greater than that in PC-9. This observation is consistent with clinical findings showing that patients harboring exon 19 deletion-type EGFR mutations often respond more favorably to therapy than those with the L858R point mutation^49^. Interestingly, certain resistance mutations were identified within regions encoding the extracellular domain of EGFR. These mutations might contribute to resistance to osimertinib, by altering ligand binding or interfering with EGFR homodimerization similar to R84K and A265 mutations^50^. Several of the resistance mutations identified in our screening overlapped with previously reported driver mutation hotspots, suggesting that these variants have clinical significance. Many of the mutations flagged as resistance-associated in this study have been rarely reported in clinical databases, and thus may represent underappreciated contributors to drug resistance. However, when aggregated across patients and treatment contexts, such mutations could represent a substantial burden to therapy efficacy (**Fig. 4h**). These findings underscore the importance of compiling comprehensive mutation-specific resistance profiles for each targeted therapy, which could serve as a valuable resource for precision oncology.

BELT enables high-throughput functional mutation screening with higher speed and scalability than previously reported experimental systems^10,11^. This method is thus especially suitable for applications such as rapid functional characterization of pathogen susceptibility variants during pandemics and the accelerated development of next-generation molecular-targeted drugs. Beyond resistance mutations, BELT-based screening may also be applicable to the systematic evaluation of variant-dependent drug responses, offering a powerful strategy for expanding the functional landscape of pharmacogenomics.

## Materials and Methods

### Cell culture

HEK293 cells were obtained from Japanese Collection of Research Bioresources Cell Bank (#9068) and KMST-6 cells were obtained from RIKEN BioResource Research Center (BRC) (#RCB1955). Cells were maintained in Dulbecco’s Modified Eagle Medium (DMEM) (#10566016; Thermo Fisher) containing 10% fetal bovine serum (FBS) and 100 μg/mL penicillin–streptomycin. II-18 cells and PC-9 cells were obtained from the RIKEN BRC (#RCB1637, #RCB4455) and maintained in RPMI medium (#11875093; Thermo Fisher) containing 10% fetal bovine serum and 100 μg/mL penicillin–streptomycin.

### Preparation of vectors

Both pCAG-CBE4max-SpRY-P2A-EGFP (RTW5133) and pCMV-T7-ABEmax(7.10)-SpRY-P2A-EGFP (RTW5025) were a gift from Benjamin Kleinstiver (Addgene plasmid #139999 and #140003). After treatment of these vectors with the restriction enzyme SapI (#R0569S; NEB), the sequence of phosphoglycerate kinase promoter -driven puromycin resistance gene (pPGK-puro) was inserted into the vectors using HiFi DNA Assembly Master Mix (E2621S; NEB) to prepare CBE-SpRY-Puro and ABE-SpRY-Puro. pX330-U6-Chimeric_BB-CBh-hSpCas9 was a gift from Feng Zhang (Addgene plasmid # 42230). For preparation of the pEX-U6 vector, sequences of the U6 promoter derived from pX330 (Addgene plasmid #42230) and pPGK-puro were inserted into pEX-A2J2 vector (Eurofin Genomics).

### Preparation of the gRNA library

PCR amplicons of CDSs of six genes were prepared using the SuperScript IV RT-PCR kit (#12595025; Thermo Fisher) and primers are listed in Supplementary Table S4. The PCR reaction solution was separated by agarose gel electrophoresis and DNA in the expected size of bands was purified using a GEL extraction kit (#28706; Qiagen). PCR amplicons were randomly digested using a dsDNA Fragmentase kit (#M0348L; NEB) to a size of 200-500 bp, and terminals were blunted using a Quick blunting kit (#E1201L; NEB). The Stem-loop DNA, Adaptor_A, the oligo sequence of which is shown in Supplementary Table S5, was ligated into the blunt end of fragmented DNA using the Quick ligation kit (#M2200L; NEB), and the fraction of ligated DNA measuring >150bp in length was purified using Select-a-Size DNA Clean & Concentrator kit (D4080; ZYMO RESEARCH). Target DNA was amplified with the KAPA HiFi PCR kit (KK2602; Nippon Genetics) with the M13 primer after cleavage of the hinge in Adaptor_A by USERII enzyme (#M5508S; NEB). Then, DNA measuring 150–500 bp size of DNA was purified by GEL extraction kit.

The fractionated DNA was treated with EcoP15I (#R0646S; NEB), and DNA <100 bp was filtered out using a Select-a-Size DNA Clean & Concentrator kit. Adaptor_B, for which sense and antisense oligo sequences are shown in Supplementary Table S5, was ligated into the 5’-protruding end of the DNA fragment with Adaptor_A using a DNA ligation kit (#6023; Takara Bio), and the assembled DNA fragment was collected with streptavidin magnetic beads (#65601; Thermo Fisher). As the PCR template of the bead-DNA complex, target DNA was amplified using a KAPA HiFi PCR kit with InsAmp primer pairs. After treatment with BamHI (#R3136T; NEB) and XhoI (R0146S; NEB), the digested PCR amplicon was separated by electrophoresis using a TBE gel (#EC6275; Thermo Fisher), and 110 bp-sized DNA bands were purified. This DNA fragment was inserted into the pEX-U6 vector as a spacer sequence and transformed into E. coli with high competency (#310-07733; NIPPON GENE). After incubation on agarose plates of LB medium with ampicillin at 37℃ overnight, vectors were collected from colonies and purified using the Plasmid Maxi Kit (#12163; Qiagen).

### Evaluation of gRNA library quality

The PCR amplicon containing the spacer sequence was prepared as a template of the gRNA library with the Insert_check_Fwd and Insert_check_Rev primers shown in Supplementary Table S5. A second PCR was performed to tag the barcode sequence at the end of the strand, and library quality was checked using the dsDNA 915 Reagent Kit (#DNF-915-K0500; Agilent). The sequence was then analyzed by Amplicon-Sequencing with NextSeq 1000 (2 × 300 bp) using the NextSeq 1000/2000 P1 Reagents kit (#20075294; Illumina). The DNA sequence encoding spacer RNA was identified according to the position of the U6 promoter and tracrRNA sequences. To evaluate the off-target effect of the gRNA, sequence lists of all possible 20-bp spacers along the target gene were prepared and the number of hits for each one against human genome (hg38) was analyzed with the GGGenome tool (https://gggenome.dbcls.jp/en/) in DBCLS.

### BELT

Each target of the gRNA library was pooled in proportion to the length of the target genomic region, and a mixture of CBE-SpRY-PURO, ABE-SpRY-PURO, and the gRNA library was prepared using equal weights. For HEK293 cells, 60 µg of vector mixture was transfected with 1.0 × 107 cells by NEON Electroporation system (#MPK5000; Thermo Fisher) with the following parameters (voltage: 1,000 V, Width: 30 ms, Pulses: 2 pulses). For the KMST6 and II-18 cell lines, similarly, cells were transfected with vectors using a Lipofectamine 3000 kit (#L3000008; Thermo Fisher). Two days after transfection, the cells were continuously cultured in medium with puromycin (1.5 μg/ml in HEK293 and II-18, 3.0 μg/ml in KMST-6) for 48 h, and then maintained in normal medium for 4–5 days for cell expansion.

### Enrichment of the cellular fraction harboring functional variants

For HEK293 or KMST-6, 1.0 × 106 cells harboring random SNVs in the CDS of six oncogenes were cultured on 100 mm ultra-low attachment dish (#4615; Corning) for 5 days. The cell culture conditions were established following the reported study for GILA assay^51^. The culture medium containing spheroids was transferred to a 15 ml tube, and the tube was left undisturbed for 1 min to collect a relatively large mass of spheroids with specific gravity. The cell pellet was suspended again in 12 ml of PBS, and left undisturbed for 1 min, and this step was repeated four times. To select the cells with drug-resistant variants, 1.0 × 106 cells of II-18 or PC-9 harboring random SNVs in the CDS of the EGFR gene were incubated on 100 mm ultra-low attachment dish overnight and treated with 0.5 μM or 0.2 μM osimertinib (#S7297-10mM; Selleck) respectively, for 10 days.

Each candidate of driver mutation was introduced into HEK293 cells using expression vectors for gRNA with the short guide RNA sequence listed in Supplementary Table S6 and BEs. The induction of target SNVs was confirmed by Sanger sequencing (Supplementary Figure 1). These lines of HEK293 cells were suspended in DMEM containing 1.0% FBS, and 500 cells/100 μl were seeded per well in an ultra-low attachment 96-well plate (#3474; Corning) (each line, n = 6). On days 0, 2, and 5 after starting culture, cell viability was quantified using CellTiter-Glo 2.0 (#G9243; Promega) following the manufacturer’s protocol.

### Preparation of amplicon DNA for target genes

Total RNA was purified from the target cell fraction using the Trizol reagent (#15596026; Thermo Fisher), and mRNA was further purified using the RNeasy kit (#74104; Qiagen). PCR amplicons of the mRNA of six genes were prepared using a SuperScript IV RT-PCR kit, and the primers used are listed in Supplementary Table S4. The fraction of pre-culture samples was parallelly treated by the same way.

### NGS analysis

For analysis with a short read sequencer, amplicons prepared from pre- or post-treatment samples were pooled in equimolar amounts according to size and digested with a dsDNA Fragmentase kit to a size of 400–500 bp. Fragmented DNA was used for library preparation using the NEBNext Ultra II DNA Library Prep Kit (#E7645; NEB) with unique-dual index adaptor from NEBNext Multiplex Oligos for Illumina Kit (#E7395; NEB). Library concentration and quality were examined by DNA ScreenTape (5067-5365; Agilent) using the 2200 TapeStation (Agilent). Paired-end sequencing (2 × 250 cycles or 2 × 150 cycles) was performed on the Illumina Miseq with Reagent kit v2 (#MS-103-1002 or MS-103-1003; Illumina) at Tsukuba i-Laboratory LLP.

For HiFi long read sequencing analysis, amplicons of EGFR mRNA were prepared with amino modifier C6-primers. After the second PCR for tagging of the barcode sequence at the end of each strand, SMRTbell libraries were prepared using barcoded universal primers for multiplexing amplicons according to the PacBio protocol. The DNA sequence of full-length EGFR mRNA was analyzed using the HiFi-mode of the Sequel IIe System (PacBio) using the Sequel II binding kit 2.1 (#101-820-500; PacBio). HiFi reads were computationally prepared using SMRT Link (v12.0.0.177059) for treatment of reads with circular consensus sequencing.

### Identification of enriched mutations

The sequence reads in FASTQ data were first trimmed using the “Trim Reads” command and the CLC genomics workbench (v25.0.1; QIAGEN) (CLC-GW), and filtered reads were mapped against the transcript’s sequence (hg38) using the “RNA-Seq Analysis” tool of CLC-GW. The numbers of counts and coverages on each variant registered in the COSMIC and ClinVar databases were calculated using the “Identify Known Mutations from Mapping” command of CLC-GW. After filtering out variants with coverage <8,000, mutation frequencies were obtained from count per coverage. For detection of DEMs, FC (ratio of mutation frequency) and p value (Fisher’s exact test for the cross-tabulation table of counts and coverage) between two groups were calculated.

For analysis of osimertinib resistance, after trimming using CLC-GW, the sequences were first aligned to the CDS of the EGFR gene, and reads containing nonsense mutations were excluded from downstream analysis to avoid the complexity of interpreting functional consequences arising from truncating forms caused by nonsense mutations. Followed by mapping, in osimertinib-treatment samples, single variants and variant pairs with counts <5 were filtered out for DEM calling. SNVs (mutation frequency >10%; without L858R mutation) present in the parental II-18 cell line prior to the introduction of mutations were omitted from the pairwise mutation analysis. The Circos plot was generated using the Circos software package (https://circos.ca)^52^.

### Genome coordinates

Genomic features including exon, intron, and coding sequences were defined with ncbiRefSeq data obtained using the UCSC table browser (hg38). The variant call file (VCF) of COSMIC variants (v98 on GRCh38) was downloaded from the COSMIC website. The VCF of ClinVar with annotation including genomic position, pathogenicity, and related disease names on six genes was obtained from Table Browser of the UCSC website (hg38).

### Comparing mutations with variant databases

The integrated variants from COSMIC and ClinVar were analyzed by the Variant Effect Predictor tool (v114) on the Ensemble website, and scores of Polyphen-2 CADD, ClinPred, AlphaMissense, and SpliceAI were obtained using this tool. The mRNA secondary structure was predicted and analyzed with the python script of SNPfold (v1.0)35. The table of codon usage in Homo sapiens was obtained from the Codon Usage Database^53^ in the webpage of Kazusa DNA Research Institute. The scores were divided into two groups according to associated variants of DEMs and non-DEMs, and the median values of each group were calculated.

### Xenograft model

The expression vectors for CBE and ABE with or without gRNA libraries were introduced into HEK293 cells. After puromycin selection for 48 h, cells were incubated on a high attachment dish (#3020-100; IWAKI) for 1 week to promote expansion. Harvested cells with or without gRNA (2.0 × 10^6^ each) were suspended in MatriMix (#899-001; Nippi), and then transplanted into the left or right shoulder of immunodeficient SCID male mice obtained from Charles River Laboratories, respectively. Tumor size was measured every other day from day 30 to day 44 after cancer cell transplantation. On day 44, the mice were euthanized, and subcutaneous tumor formation was confirmed. Xenograft tumors were fixed with PBS containing 4% paraformaldehyde and embedded in paraffin. The successive sections were stained using hematoxylin and eosin, and the expressions of proliferation markers were detected by DAB staining kit (#414341 [for Ki67] or #414171 [for PCNA]; NICHIREI BIOSCIENCES) with anti-Ki67antibody (ab16667; Abcam) and anti-PCNA antibody (#2586; CST).

### Drug resistance assay

Each candidate drug-resistant mutation was introduced into II-18 cells by expression vectors for both gRNA with a short guide RNA sequence listed in Supplementary Table S6 and BE. The induction of target mutations was confirmed by Sanger sequencing (Supplementary Figure 1). II-18 cells harboring SNVs were suspended in RPMI containing 10% FBS, and 4.0 × 10^3^ cells/100 μl were seeded per well in an ultra-low attachment 96-well plate. After overnight incubation to induce spheroid formation, medium containing osimertinib was added into each well, adjusting to each concentration (each concentration n = 6). On day 3, cell viability was quantified using CellTiter-Glo 2.0. Preparation of dose-response curves and evaluation of the differences in the drug response were achieved by R package “DRC” ^54^.

### Western blotting

II-18 (3.0 × 10^5^ cells) with each SNV were cultured on ultra-low attachment 60 mm dishes (#3261; Corning). After overnight culture, osimertinib was added into the dishes at a final concentration of 100 nM. After treatment for 2 h, the collected spheroids were immediately suspended in 1 × SDS sample buffer and boiled for protein denaturation after sonication to shear genomic DNA. The blotted membranes were incubated with the primary antibodies listed in Supplementary Table S7, and signals were detected using a peroxidase-conjugated anti-mouse secondary antibody (#G21040; Thermo Fisher) or anti-rabbit secondary antibody (#G21234; Thermo Fisher), followed by treatment with ECL + reagent (#34075; Thermo Fisher). Membrane images were scanned by FUSION FX imaging systems (Vilber Bio Imaging) (Supplementary Figure 2).

### Quantification and statistical analysis

In all box plots, elements indicate the following values: center line, median; box limits, upper and lower quartiles 1.5× interquartile range; and points, outliers, and median values of each group were compared by two-sided Wilcoxon rank-sum test. For differences in mutation frequency, p values were calculated by Fisher’s exact test for numbers of read counts and coverage in each variant and then converted into FDR. In the spheroid formation assay, the differences in cell growth on day 5 between the wild type and mutants were evaluated using Welch’s test. In the dose-response curve, the differences were evaluated using the “comped” function of R package “drc”^54^. The sample size in this study was decided based on past experience in generating statistical significance. Investigators were not blinded to experimental conditions, and no randomization or exclusion of data points were used.

## Data availability

All sequencing data generated using the next-generation sequencer in this study were deposited in the NCBI Sequence Read Archive (SRA) under the BioProject number, PRJNA1274207.

## Acknowledgments

We would like to thank i-Laboratory staff in Tsukuba university for supporting the NGS analysis and Dr. Kurisaki in Research Institute for Science and Engineering, Waseda University for comments on the 3D structure of the EGFR protein. This research was supported by Research Support Project for Life Science and Drug Discovery (Basis for Supporting Innovative Drug Discovery and Life Science Research [BINDS]) from AMED under Grant Number JP24ama121055, and the Science Research Promotion Fund of the Promotion and Mutual Aid Corporation for Private Schools of Japan.

## Author contributions

K.Y. conceived the idea underlying the principle of the BELT assay and performed variant screening experiments and cellular function assays. K.Y. and T.A. prepared gRNA expression vector libraries. N.M. performed the experiments related to the xenograft model. E.S., M.M., M.H., and S.S analyzed amplicon-sequencing data. K.H. and K.Y. evaluated the clinical significance of each variant. K.Y. and K.H. wrote the manuscript.

## Ethics declarations

### Ethics approval

The experiments were conducted in accordance with the guidelines of the Institutional Animal Care and Use Committee of Nippon Medical School.

## Additional information

M.M. is associated with Tsukuba i-Laboratory as a technical consultant. K.Y., K.H., T.A. filed a patent application related to this work and it is currently under review. Details remain confidential at this stage.

---

**Supplementary Figure 1: Original Sanger sequencing data for confirmation of SNV knock-in.**

**Supplementary Figure 2: Uncropped image of western blot for protein expression.**

**Supplementary Table S1: List of COSMIC mutations with FC and FDR, for driver mutation screening using HEK293 and KMST-6.**

**Supplementary Table S2: List of all SNVs, 1-bp resolution, with FC and FDR, for driver mutation screening using HEK293.**

**Supplementary Table S3: List of COSMIC mutations with FC and FDR, for osimertinib-resistant mutation screening using II-18 and PC-9.**

