## Extended Data (Extended Data Figure 1-7 & Table 1,2) for "Comprehensive and accelerated mapping of driver mutations through single-nucleotide random mutagenesis of target genes"

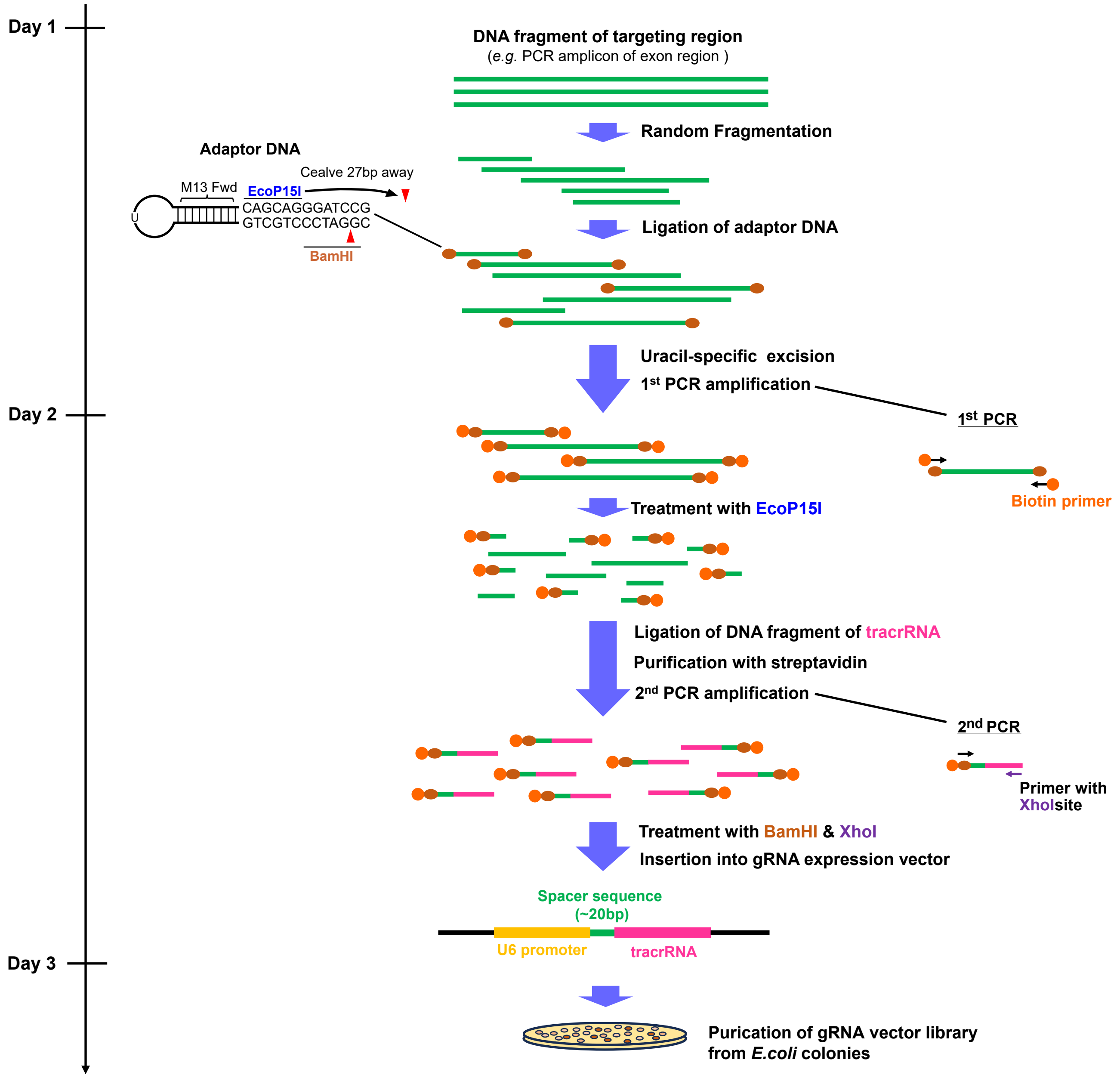


**Extended Data Fig. 1 A step-by-step protocol for gRNA library preparation.**

To produce variety of terminal sequences, DNA fragment of target region (green line) was sheared by fragmentase. After blunting ends, Adaptor_A stem-loop DNA (brown oval) which contains recognition sites of EcoP15I and BamHI, is ligated into terminals of target, and it is amplified by biotin-tagged M13_Fwd primer specifically to produce DNA fragment flanked by adaptor_A on both ends with biotin (orange circle). Treatment with EcoP15I induces cleavage 20bp away from ligation site, and Adaptor_B, oligo DNA involving tracrRNA and XhoI recognition sequences is ligated into the fragment at opposite end of Adaptor_A, followed by the 2$^{nd}$ PCR amplification. After BamHI and XhoI treatment, purified DNA for “20bp + tracrRNA sequence” is inserted into gRNA expression vector.

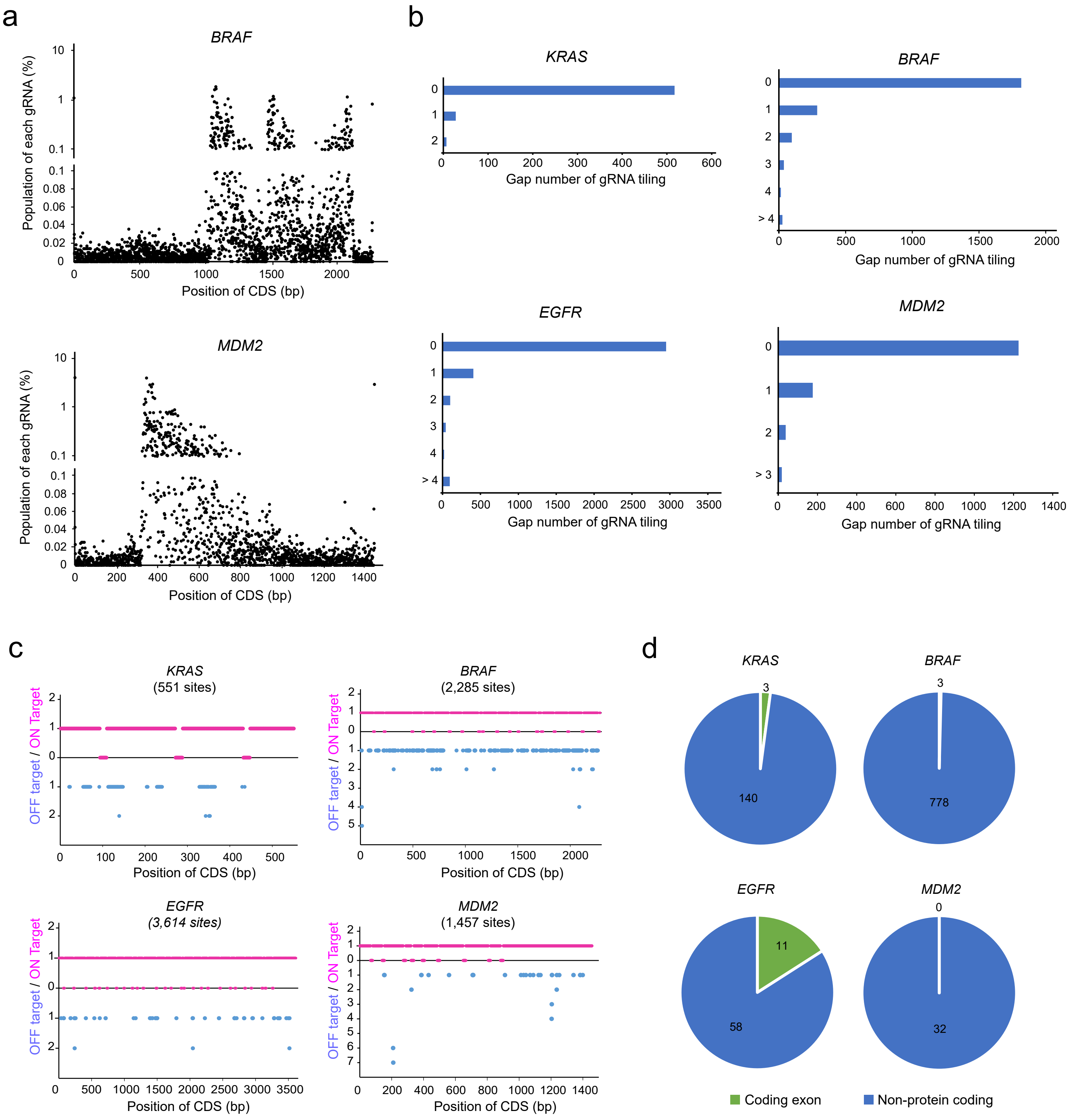


**Extended Data Fig. 2 Distribution of gRNAs on target region and assessment of off-target effect.**

**a**, Proportion of gRNA in which spacer sequence corresponds to 20bp along exon sequence of *BRAF* (top) or *MDM2* (bottom). **b**, Distribution of gap number of gRNA tiling. The gap length between each arrayed gRNA was analyzed. **c**, Simulated distributions of hit number of expected gRNA for on-target and off-target. List of all possible 20bp as spacer sequence from target CDS was prepared, and number of hits on human genomic sequence was analyzed by gggenome, high-speed sequence search tool specialized for short read. **d**, Distribution of spacer sequence of off-target gRNA against coding (exon) and non-coding region (intron, UTR, intergenic).

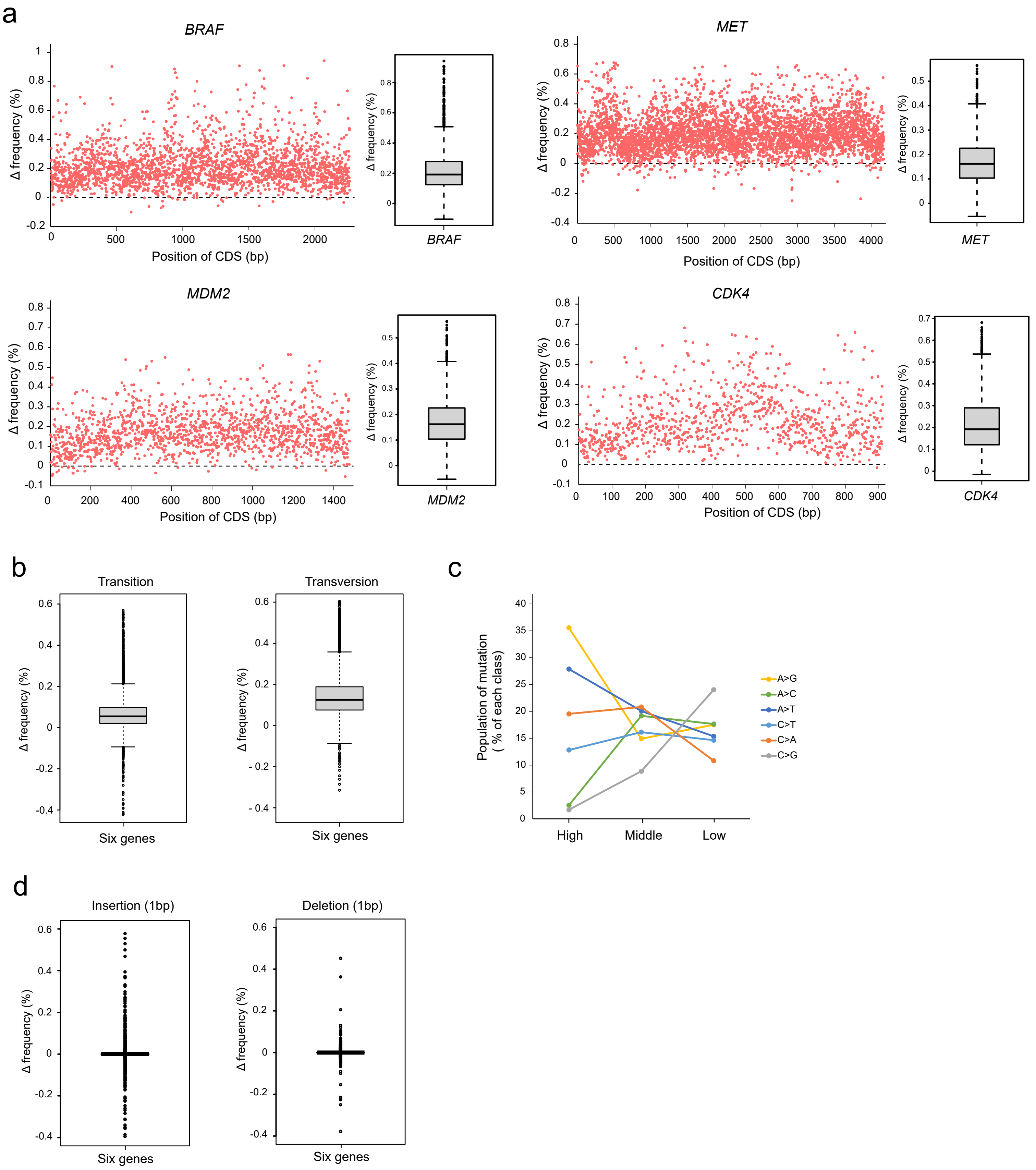


**Extended Data Fig. 3 Distribution of induced SNVs by BELT mutagenesis.**

**a**, Induction of frequency of SNV on all base of *BRAF*, *MDM2*, *CDK4*, *MET* gene. Distribution of frequencies are shown in right part of each gene (n=2,277 in *BRAF*, n= 1,476 in *MDM2*, n=912 in *CDK4*, n=4,173 in *MET*) by box plot. **b**, Distribution of frequencies of transition (n=13,041) and transversion (n=26,082) mutations on combined exon region for six oncogenes. **c**, Population of each class of mutation alterations. Mutations were classified by high (Δfrequency >0.2%, n=1,474), middle (0.2%> Δfrequency >0.05%, n=20,214), and low (0.05%> Δfrequency >0.01%, n=5,995). **d**, Distribution of frequencies of 1bp-insertion (n=13,041) and 1bp-deletion (n=13,041) on combined exon region for six oncogenes.

Yoshida K et al., Extended Data Figure 4

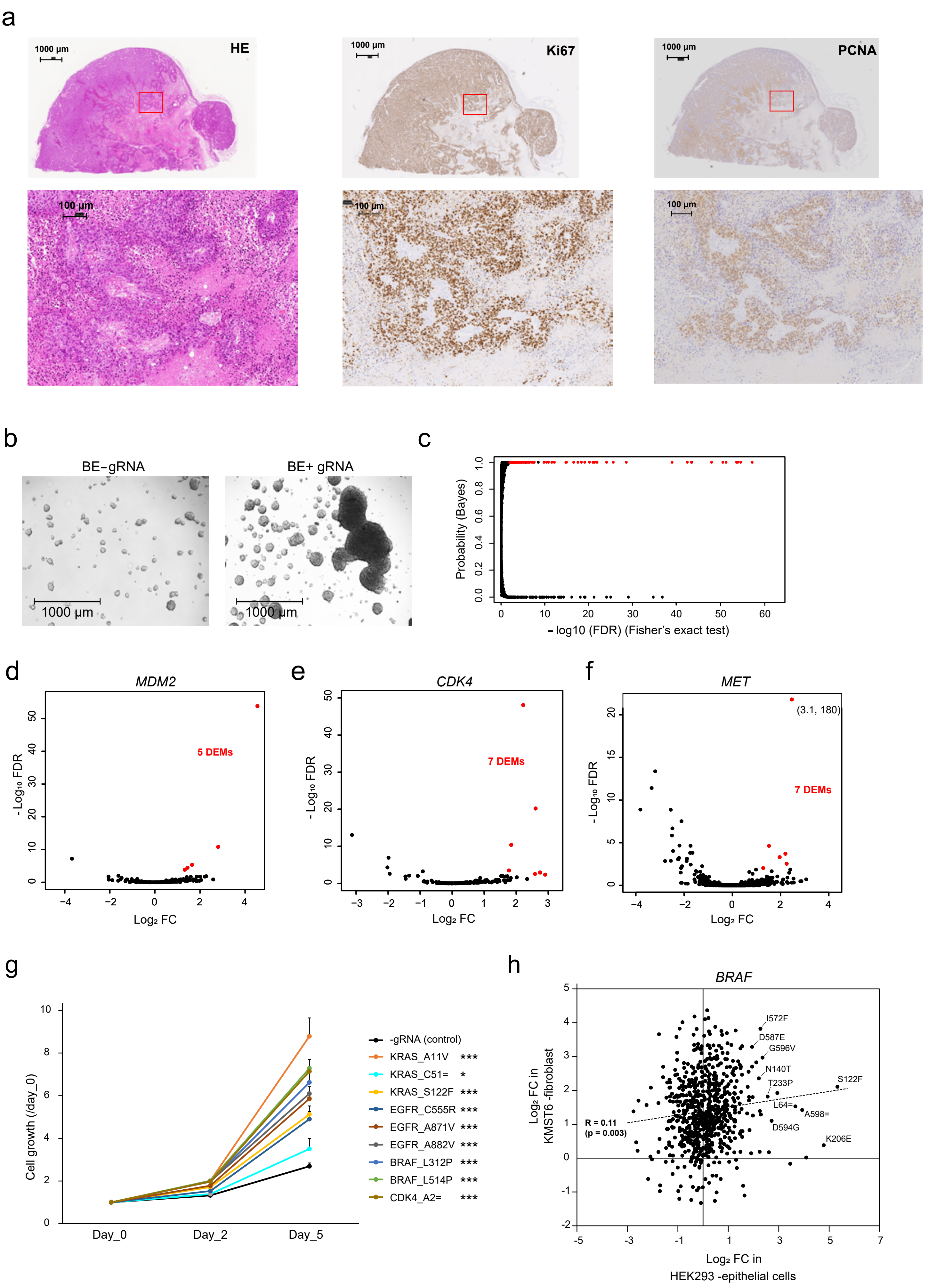

**Extended Data Fig. 4 Driver mutations are enriched in the genome of cultured cells that have acquired tumorigenic potential.**

**a**, Representative image of HE staining (left) and staining with Ki67 (middle) or PCNA (right) antibody in the successive sections of tumors formed in vivo. The enlarged images were shown in the below part. **b**, Representative image of spheroid from HEK293 treated with only BE or both BE and gRNA library, formed on low attachment dish under bright-field microscopy. **c**, Relationship between p value calculated by Fisher's exact test and probability calculated by Bayesian inference, for evaluation of the difference of mutation frequency in post-sample relative to pre-sample. **d-f**, Volcano plot for COSMIC variants on *MDM2* (**D**), *CDK4* (**E**), and *MET* (**F**) in HEK293. DEMs (FC >2, FDR <0.01) are colored by red, and representative mutations are shown with amino acid alteration. The number of data points falling outside the plotting range is annotated at the plot margins. **g**, Result of spheroid formation assay. Cells harboring each COSMIC-registered point mutation were cultured on low binding dish for 5days. For quantification of anchorage-independent growth ability, cell viabilities were measured at day0, 2, 5 (n=6 at each time point), and they were normalized by day0. Difference of the ability was evaluated by relative viability of control to each mutant line at day5, by Welch's test. **h**, Scatter plot for FC in current our results of *BRAF* gene between epithelial-like and fibroblast-like cell-lines. *P<0.05, ***p<0.001.

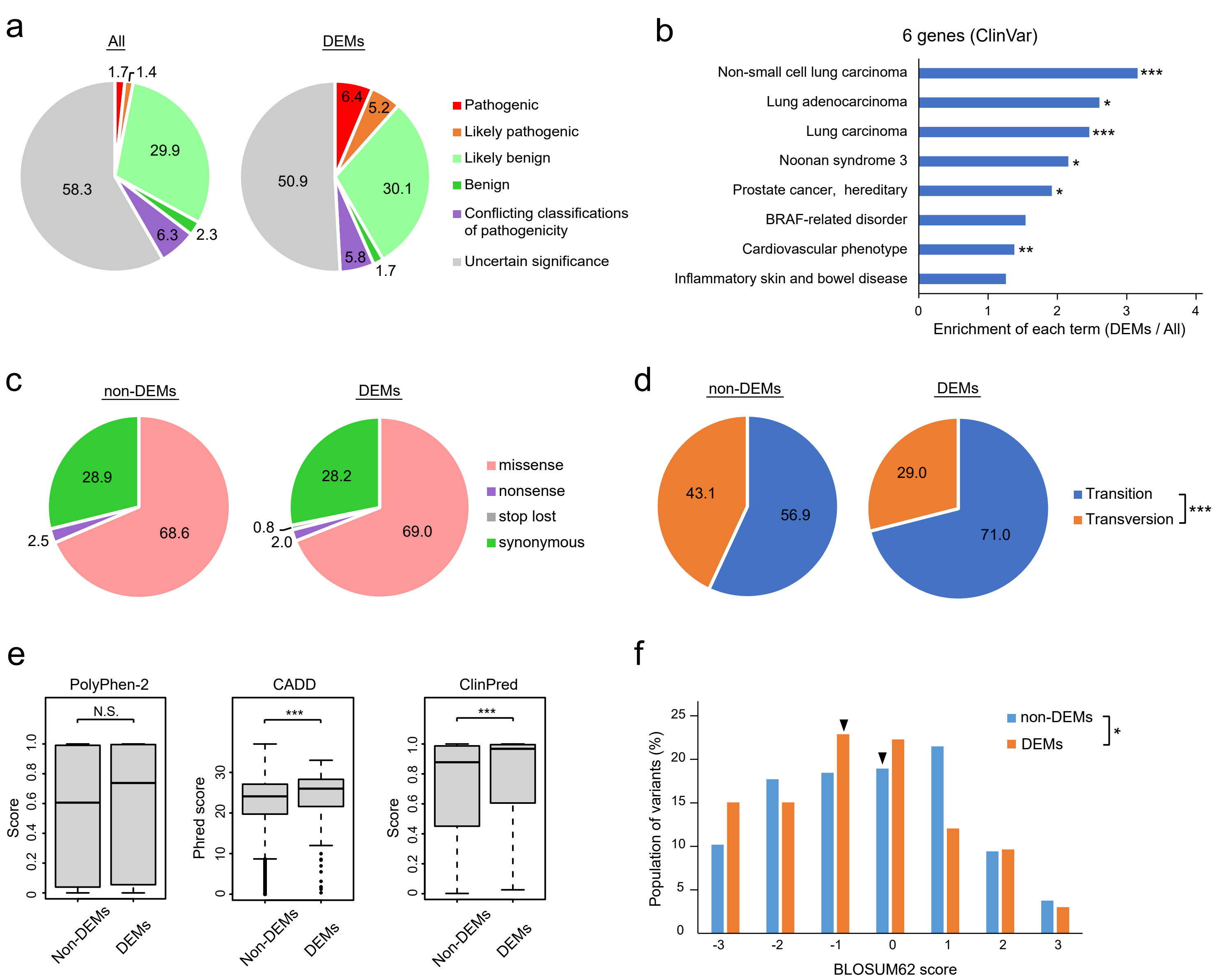


**Extended Data Fig. 5 Fraction of each mutation property in DEMs.**

**a**, Pie chart for population of ClinVar variants with annotations of clinical significance. **b**, Enrichment of the variant population with annotations of related disease in ClinVar between DEMs (FC >2, FDR <0.01) and all mutations. P value was calculated by Fisher's exact test. The top eight disease terms are shown. **c** and **d**, Pie chart for population of combined list of COSMIC and ClinVar variants classified by mutation classification and base substitution type. **e**, Distribution of PolyPhen, CADD or ClinPred scores of missense variants in COSMIC and ClinVar databases between non-DEMs (n = 6,810) and DEMs (n = 166). P value was calculated by Wilcoxon rank-sum test. **f**, Population of BLOSUM62 scores of missense variants in COSMIC and ClinVar databases. The difference of population in each score was evaluated by Fisher's exact test. Median score of each group was indicated by black triangle. N.S., not significant, *P<0.05, ***p<0.001.

Yoshida K et al., Extended Data Figure 6

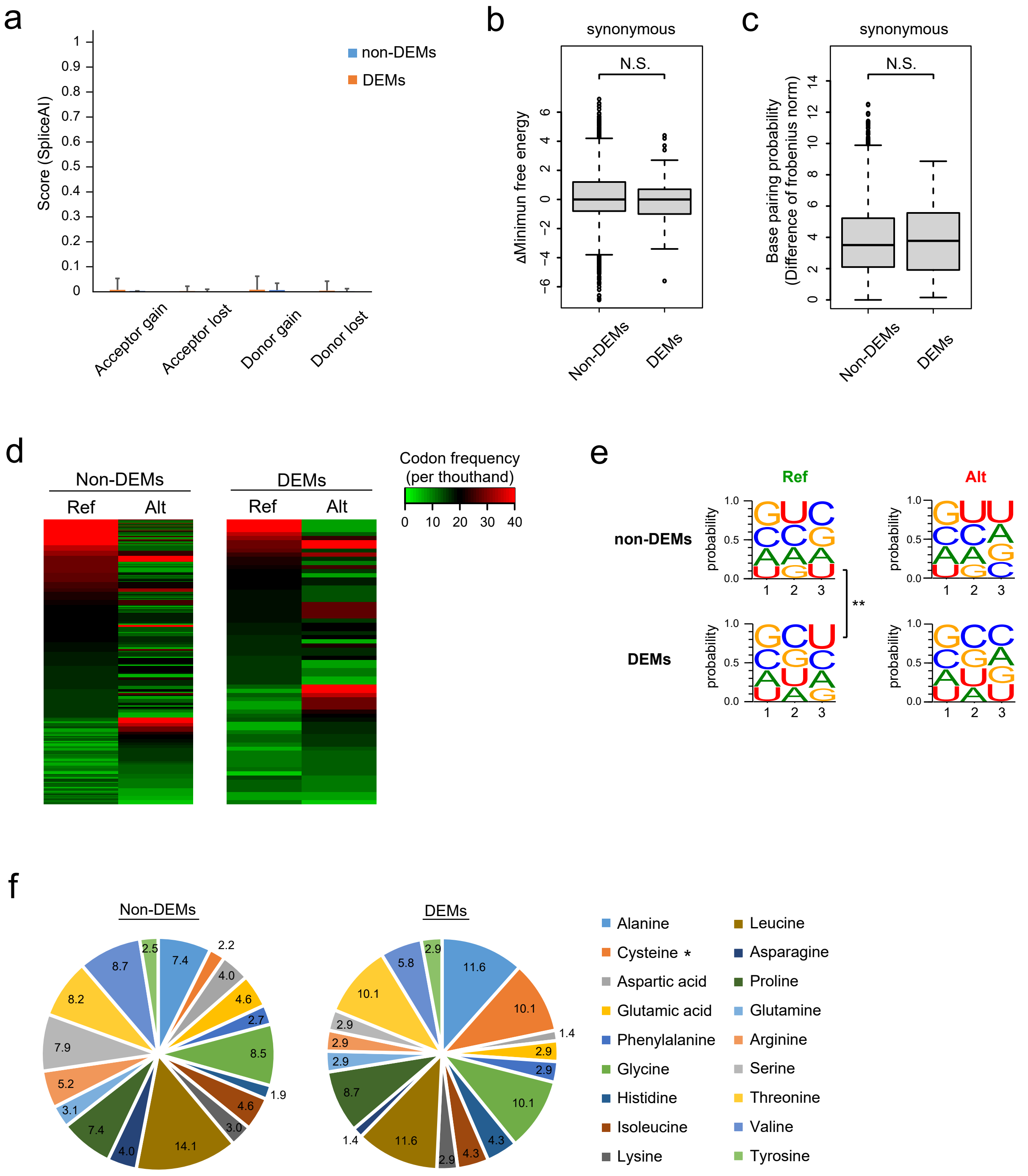


**Extended Data Fig. 6 Evaluation of the effect of synonymous DEMs on mRNA properties.**

**a**, Scores of SpliceAI for each class for all synonymous mutations between non-DEMs and DEMs. **b** and **c**, Distribution of Δ minimum free energy and base paring probability of synonymous variants in COSMIC and ClinVar databases between non-DEMs and DEMs. P value was calculated by Wilcoxon rank-sum test. **d**, Heatmap of usage frequency in the reference and alteration codon between non-DEMs and DEMs. **e**, Nucleotide probabilities in the reference and alteration codons. **f**, Pie chart for coding codons of each amino acid between non-DEMs (n = 2,792) and DEMs (n = 69). To estimate the difference in nucleotide appearance, P values are calculated by Fisher's exact test.

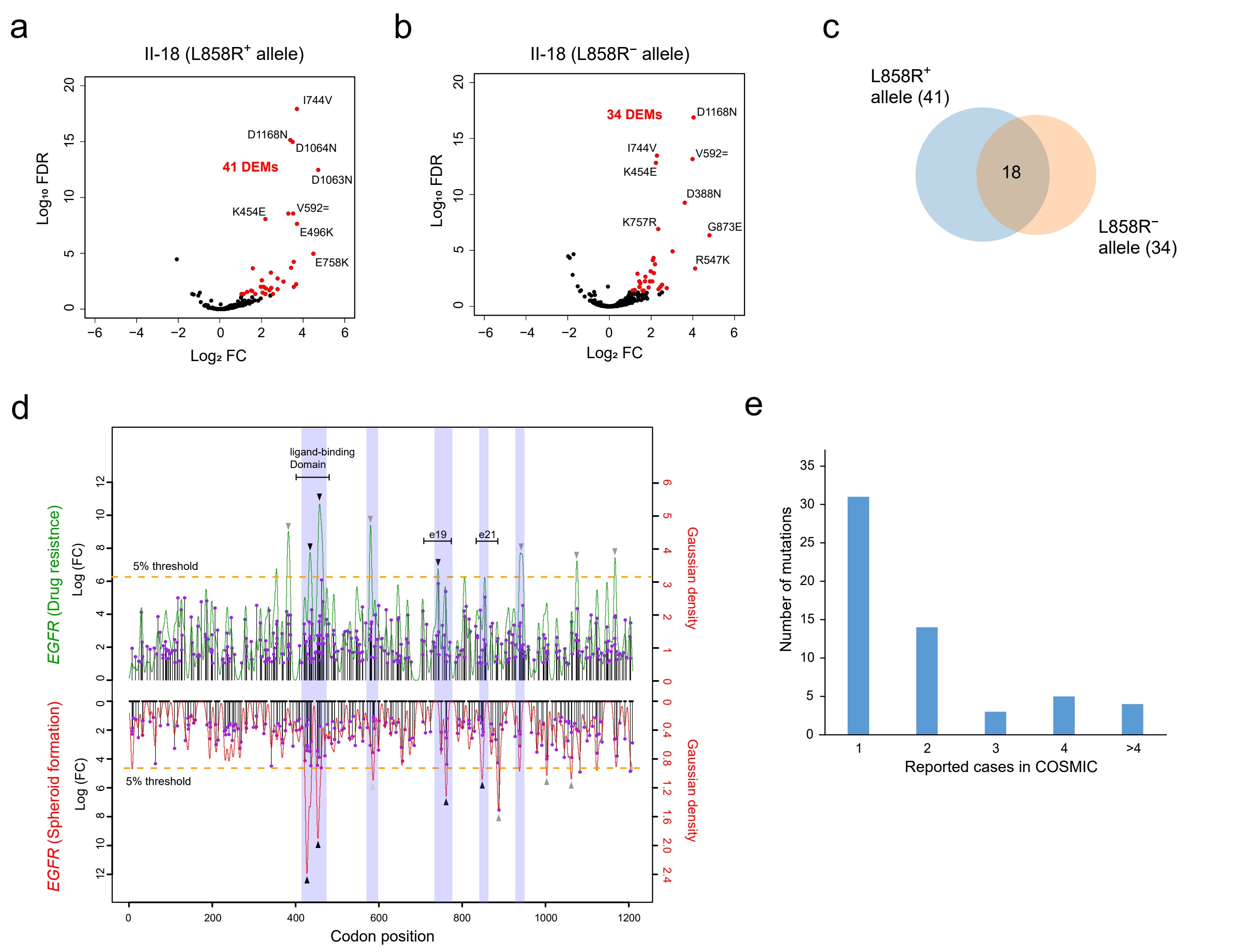


**Extended Data Fig. 7 Functional relationship of driver mutation and single nucleotide-level distributions of drug-resistant mutations.**

**a** and **b**, Volcano plot for COSMIC variants on *EGFR* gene allele with L858R and without L858R mutation in II-18. DEMs (FC >2, FDR <0.05) are colored in red, and representative mutations are shown with amino acid alteration. P values were calculated by Fisher's exact test. The number of data points falling outside the plotting range is annotated at the plot margins. **c**, Venn graph of DEMs between *EGFR* gene allele with L858R and without L858R. **d**, Lolipop plot of DEMs (FC>2, FDR<0.05) with FC for osimertinib-resistance (top) and spheroid formation-related (bottom) mutations at 1bp resolution along *EGFR* gene. Gaussian density curve of FC is plotted in red line, and top 5% region of the density is shown by orange dot line. Hot spots of density of DEMs over 5% threshold are indicated by gray triangle, and overlapping region between drug-resistance and spheroid formation-relating were colored by blue rectangle. **e**, Histogram of the number of variants corresponding to the reported case counts in COSMIC database.

**Extended Data Table 1 Database records for mutations subjected to functional analysis.**
N.R., not registered

| | Mutation type | COSMIC_ID | Annotation in OncoKB | ClinVar | AlphaMissense class |
|---|---|---|---|---|---|
| KRAS_A11V | missense_variant | COSM511 | N.R. | N.R. | likely_pathogenic |
| KRAS_C51= | synonymous_variant | COSM5009216 | N.R. | N.R. | - |
| KRAS_S122F | missense_variant | COSM6977243 | N.R. | N.R. | likely_benign |
| EGFR_C555R | missense_variant | COSM6214981 | N.R. | N.R. | likely_pathogenic |
| EGFR_A871V | missense_variant | COSM710350 | Inconclusive | N.R. | likely_pathogenic |
| EGFR_A882V | missense_variant | COSM9520897 | N.R. | N.R. | likely_pathogenic |
| BRAF_L312P | missense_variant | COSM6946221 | N.R. | N.R. | likely_benign |
| BRAF_L514P | missense_variant | COSM6176428 | N.R. | N.R. | likely_pathogenic |
| CDK4_A2= | synonymous_variant | COSM1578969 | N.R. | N.R. | - |

**Extended Data Table 2 Characteristics of the BELT method and previous large-scale mutational screening approaches.**

| | Vector components of the system | Cost of constructing a gRNA vector library | Time required to construct a gRNA vector library | Diversity of gRNAs in the library | Design of specific guide RNAs |
|---|---|---|---|---|---|
| **BELT system** | Base Editor<br>+<br>gRNA vector library<br>(*E.coli*) | < $200<br>( /library) | For 3 days | Approximately equal to the number of bases in the target gene region | Not required |
| **Previously reported system** | Base Editor<br>+<br>gRNA vector library<br>(Lenti virus) | $10,000–$50,000<br>( /library) | For a few months | 2,000-30,000 species | Technical expertise and appropriate design tools |
