## Supplementary data (Supplementary Figure 1,2 & Supplementary Table 4-7) for "Comprehensive and accelerated mapping of driver mutations through single-nucleotide random mutagenesis of target genes"

Yoshida K et al., Supplementary Figure 1

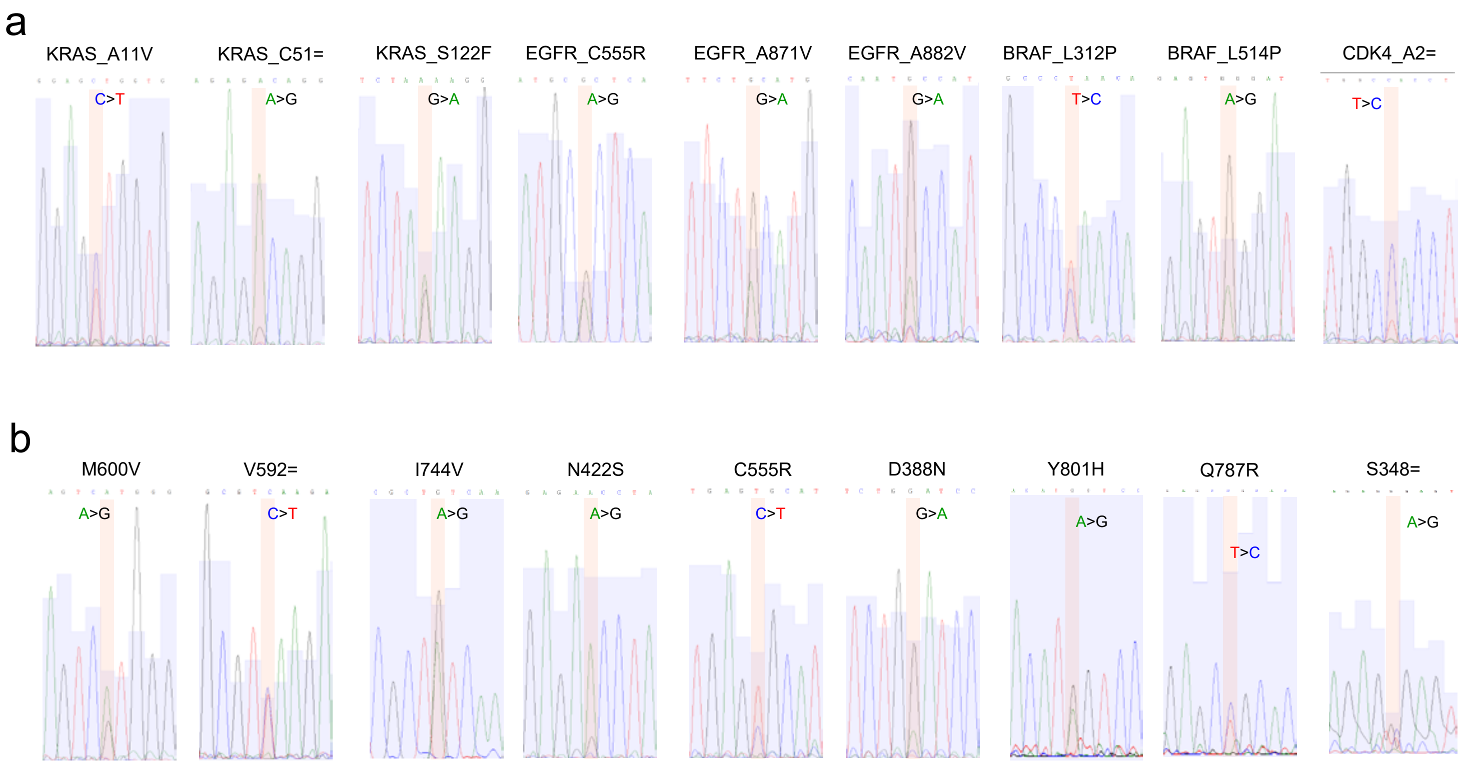


**Supplementary Figure 1 | Original Sanger-sequencing data for confirmation of SNV knock-in.**
(**a** and **b**) A trace data of Sanger-Sequencing for PCR amplicon of genome DNA collected from HEK293 (**a**) and II-18 (**b**) for checking SNV induction.

a

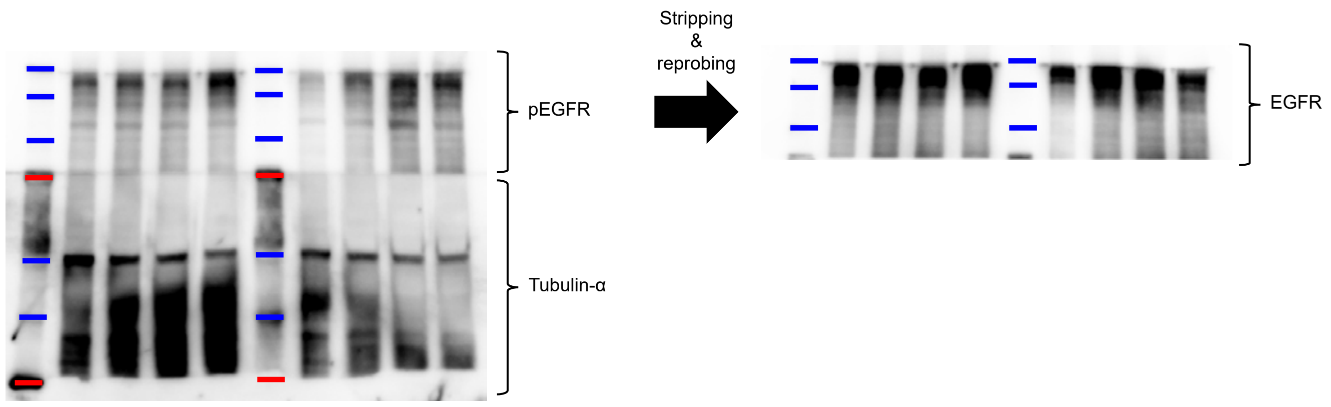


b

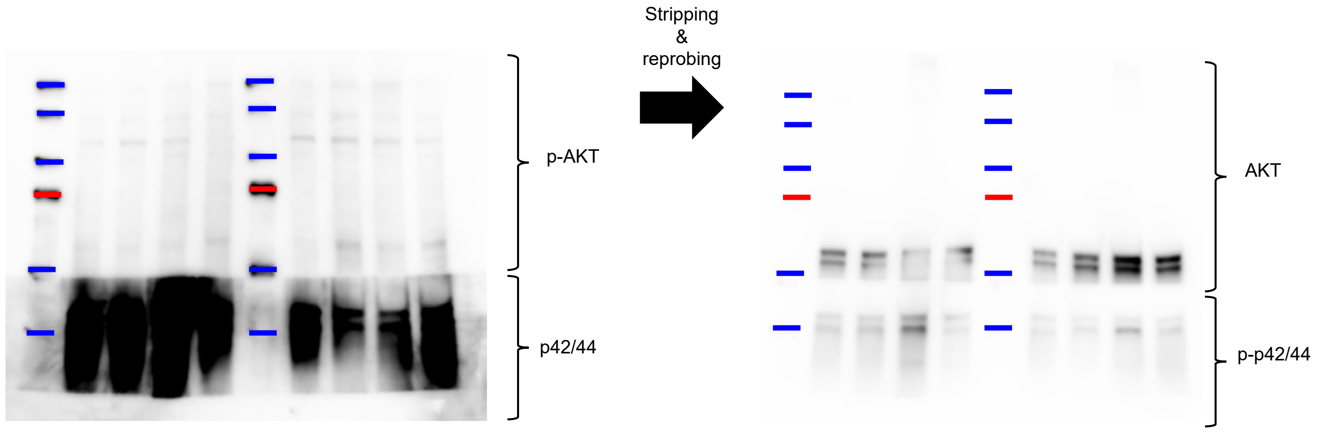


**Supplementary Figure 2 | Uncropped image of western blot for protein expression**

(**a** and **b**) Original western blot image for expressions of EGFR & Tubulin-α (**a**) and ERK & AKT (**b**).

**Supplementary Table 4 | RT-qPCR primers for oncogene mRNA**

| Target region | Fwd Sequence | Rev Sequence | Note |
|---|---|---|---|
| KRAS_CDS | ATGACTGAATATAAACTTGTGGTAG | TTACATTATAATGCATTTTTTAATTTTCAC | For gRNA Library |
| EGFR_CDS | ATGCGACCCTCCGGGAC | TCACTGTGTCTGCAAATCTGCC | For gRNA Library |
| BRAF_CDS | ATGGCGGCGCTGAGCGG | AGTGGACAGGAAACGCACCATATC | For gRNA Library |
| MDM2_CDS | ATGTGCAATACCAACATGTCTGTAC | CTAGGGGAAATAAGTTAGCACAATC | For gRNA Library |
| CDK4_CDS | ATGGCTACCTCTCGATATGAG | TCACTCCGGATTACCTTCATC | For gRNA Library |
| MET_CDS | ATGAAGGCCCCCGCTGT | TGATGTCTCCCAGAAGGAGGC | For gRNA Library |
| KRAS_mRNA | GGAGAGAGGCCTGCTGAAAATG | ACAGTCTGCATGGAGCAGGA | For Ampl-Seq |
| EGFR_mRNA | GTCCAGTATTGATCGGGAGAGC | GGCTCATACTATCCTCCGTGGTC | For Ampl-Seq |
| BRAF_mRNA | CCCCGGCTCTCGGTTATAAG | ATATCCCCCTGCCTGGATG | For Ampl-Seq |
| MDM2_mRNA | GCGATTGGAGGGTAGACCTG | GGAGTTGGTGTAAAGGATGAGC | For Ampl-Seq |
| CDK4_mRNA | TGAGGGTCTCCCTTGATCTG | CATGGCAGCCACTCCATTG | For Ampl-Seq |
| MET_mRNA | TGGGCACCGAAAGATAAACC | GGACAAAGTGTGGACTGTTGC | For Ampl-Seq |

**Supplementary Table 5 | DNA sequence of adaptor and primers for gRNA library preparation**

| Oligo Name | Sequence |
|---|---|
| Adaptor_A | CGGATCCCTGCTGACTGGCCGTCGTTTTACGTCGTATCCAGTGCAGGG**U**CCG AGGTATTCGCACTGGATACGACGTAAAACGACGGCCAGTCAGCAGGGATCCG |
| Adaptor_B_S | NNGTTTAAGAGCTATGCTGGAAACAGCATAGCAAGTTTAAATAAGGCTAGTCCG TTATCAACTTGAAAAAGTGGCACCGAGTCGGTGCTTTTTTCTCGAGGTCATAGC TGTTTCCTGCCCC |
| Adaptor_B_as | CAGGAAACAGCTATGACCTCGAGAAAAAAGCACCGACTCGGTGCCACTTTTTC AAGTTGATAACGGACTAGCCTTATTTAAACTTGCTATGCTGTTTCCAGCATAGC TCTTAAAC |
| M13 primer | biotin-CGTAAAACGACGGCCAGTC |
| InsAmp_F | CGTAAAACGACGGCCAGTC |
| InsAmp_R | AGGAAACAGCTATGACCTCGAG |
| Insert_check_Fwd | ACACTCTTTCCCTACACGACGCTCTTCCGATCTCTTGAAAGTATTTCGATTTCT TGGC |
| Insert_check_Rev | GTGACTGGAGTTCAGACGTGTGCTCTTCCGATCTAGAGGGAGTGGCCAACTC |

*underlined sequences are adaptor for the 2nd PCR

**Supplementary Table 6 | sgRNA sequences for inducing SNVs by base editor**

| Target gene | CDS alteration | AA alteration | sgRNA sequence (20bp) | BE type |
|---|---|---|---|---|
| KRAS | c.32C>T | A11V | TGGAGCTGGTGGCGTAGGCA | CBE |
| KRAS | c.153T>C | C51= | AAGAGACAGGTTTCTCCATC | ABE |
| KRAS | c.365C>T | S122F | GCCTTCTAGAACAGTAGACA | CBE |
| EGFR | c.1663T>C | C555R | TGCACTCAGAGTTCTCCACA | ABE |
| EGFR | c.2612C>T | A871V | CCATGCAGAAGGAGGCAAAG | CBE |
| EGFR | c.2645C>T | A882V | GATGGCATTGGAATCAATTT | CBE |
| BRAF | c.935T>C | L312P | GATGTTAGGGCAGTCTCTGC | ABE |
| BRAF | c.1541T>C | L514P | AGAGTAGGATATTCACATGT | ABE |
| CDK4 | c.6T>C | A2= | GAGGTAGCCATTCTCAGATC | ABE |
| EGFR | c.1798A>G | M600V | GGAGTCATGGGAGAAAACAA | ABE |
| EGFR | c.1776C>T | V592= | CTGCGTCAAGACCTGCCCGG | CBE |
| EGFR | c.2230A>G | I744V | GTCGCTATCAAGGAATTAAG | ABE |
| EGFR | c.1265A>G | N422S | GAGAACCTAGAAATCATACG | ABE |
| EGFR | c.1162G>A | D388N | CTGTGGATCCAGAGGAGGAG | CBE |

**Supplementary Table 7 | List of primary antibodies used in western blotting**

| Antigen | Host species | Catalogue |
| --- | --- | --- |
| EGFR | Rabbit | #4267; CST |
| Phospho-EGFR (Tyr1068) | Rabbit | #3777; CST |
| ERK | Rabbit | #9102; CST |
| Phospho-ERK | Rabbit | #9101; CST |
| AKT | Rabbit | #4691; CST |
| Phospho-AKT | Rabbit | #4058; CST |
| Tubulin-α | Mouse | #05-829; Merck |
